# Interplay of kinetochores and catalysts drives rapid assembly of the mitotic checkpoint complex

**DOI:** 10.1101/2024.06.09.598118

**Authors:** Suruchi Sethi, Valentina Piano, Sabrina Ghetti, Verena Cmentowski, Patricia Stege, Andrea Musacchio

## Abstract

The spindle assembly checkpoint (SAC) makes mitotic exit contingent on completion of sister chromatid biorientation, but how this coordination is achieved in practice remains poorly understood. Kinetochores, megadalton chromosome attachment sites to spindle microtubules, contribute to SAC signaling. However, it is unclear whether kinetochores are mere docking sites for SAC proteins, or further contribute to co-orientation of SAC catalysts, including MAD1:MAD2 and BUB1:BUB3, to facilitate SAC signaling. Here, we combined biochemical reconstitutions of kinetochores and the SAC to address this question *in vitro*. We engineered recombinant kinetochore particles that recruit most SAC components and activate SAC signaling after induction with Rapamycin, and challenged them with a battery of impairing mutants. At approximately physiological concentrations of SAC catalysts, kinetochores were crucially required for rapid assembly of the mitotic checkpoint complex (MCC), the SAC effector. Our observations depict kinetochores as a cradle that catalyzes rapid MCC assembly by concentrating and co-orienting distinct SAC catalysts.

## Introduction

In eukaryotes, faithful chromosome segregation during mitosis depends on the formation of bioriented attachments between the sister chromatids and the mitotic spindle. The attachment of sister chromatids to microtubules is mediated by kinetochores, multisubunit microtubule-binding machines built on specialized chromosome loci called centromeres (Cheeseman, 2014; Musacchio and Desai, 2017). The microtubule attachment process is monitored by specialized machinery, also residing at mitotic kinetochores, that corrects improper kinetochore-microtubule attachments while coordinating the progression of chromosome biorientation with mitotic exit. The feedback mechanism responsible for this coordination is named the spindle assembly checkpoint (SAC) (Fischer, 2023; Lara-Gonzalez et al., 2021b; McAinsh and Kops, 2023; Musacchio, 2015). It imposes a delay to mitotic exit in the presence of conditions that interfere with the completion of biorientation.

The SAC effector, known as the mitotic checkpoint complex (MCC), is a complex of the four proteins CDC20, MAD2, BUBR1, and BUB3 (Fraschini et al., 2001; Hardwick et al., 2000; Sudakin et al., 2001). It assembles at unattached kinetochores and inhibits cell cycle progression by binding and inhibiting the anaphase promoting complex or cyclosome (APC/C) (Figure 1A) (Alfieri et al., 2016; Fang, 2002; Fang et al., 1998; Millband and Hardwick, 2002; Sudakin et al., 2001; Tang et al., 2001; Yamaguchi et al., 2016). How kinetochores control the rate of SAC assembly is a question of great interest. *In vitro*, MCC assembly from its subunits occurs spontaneously at concentrations resembling cellular concentrations, but the rate of spontaneous MCC assembly is very slow (Faesen et al., 2017; Kulukian et al., 2009; Simonetta et al., 2009). Rate-limiting is a topological rearrangement of the HORMA domain protein MAD2, whose metastable open topology (O-MAD2) needs to convert into the more stable closed topology (C-MAD2). This conversion occurs simultaneously with the binding to a “closure motif” (also known as MIM, for MAD2-interacting motif) of CDC20 (De Antoni et al., 2005; Faesen et al., 2017; Lara-Gonzalez et al., 2021a; Luo et al., 2002; Luo et al., 2004; Piano et al., 2021; Simonetta et al., 2009; Sironi et al., 2002; Yu et al., 2024).

**Figure 1.**
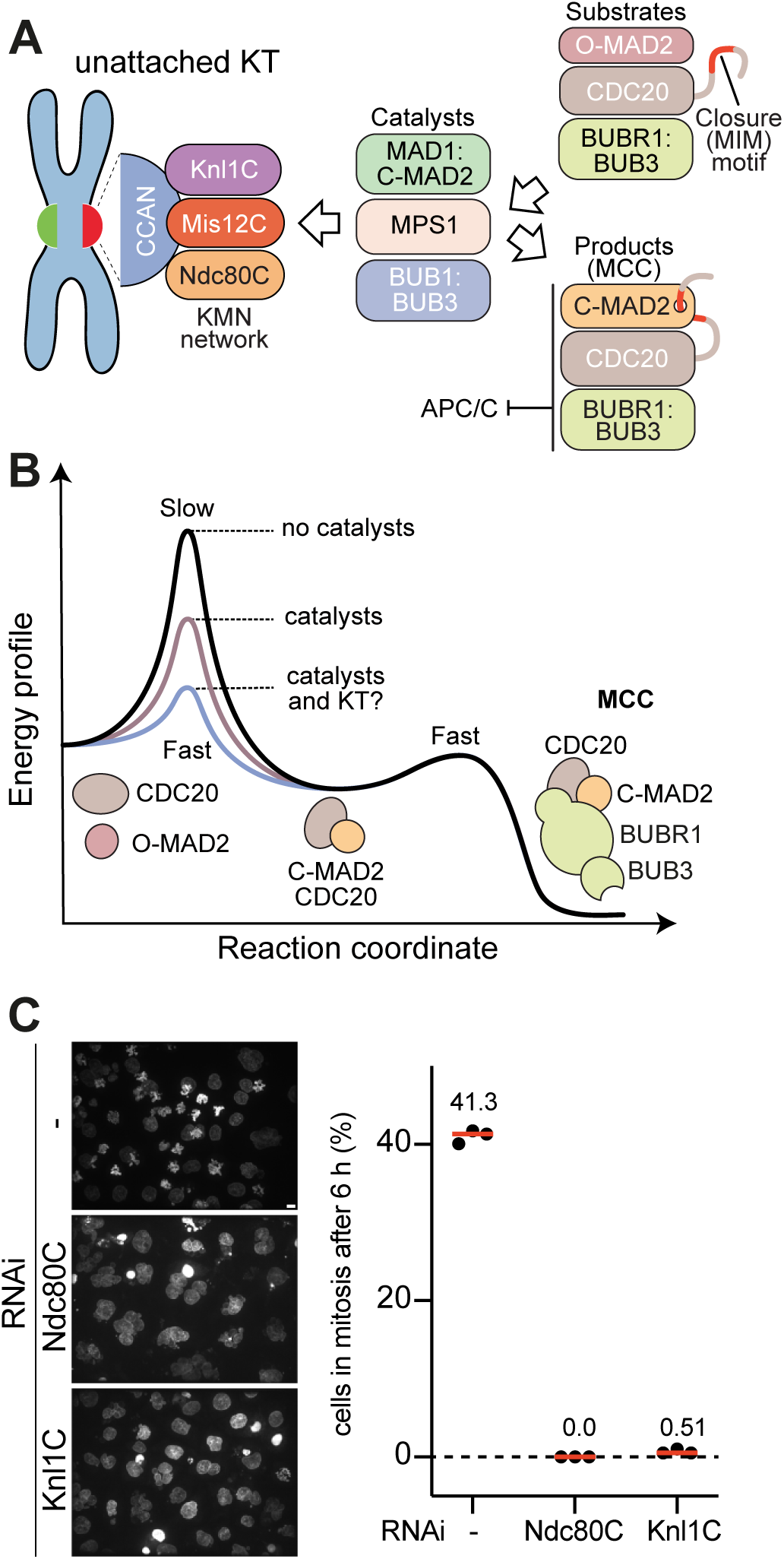
The outer kinetochore and catalysts promote SAC signaling. **A)** Scheme of catalytic assembly of MCC. Catalysts for MCC assembly are recruited to the outer kinetochore, consisting of Knl1C, Mis12C, and Ndc80C. The catalysts recruit the MCC substrates and turn them into the product, the MCC, which diffuses away and binds and inhibits the APC/C. **B)** Energy profile of MCC formation. Rate-limiting for MCC assembly is the association of CDC20:C-MAD2 from CDC20 and O-MAD2. It is spontaneous but slow due to the high activation energy. A bipartite assembly comprising BUB1:BUB3 and the MAD1:C-MAD2 core complex, activated by MPS1, catalyzes this association by lowering the energy barrier. MCC further incorporates BUBR1:BUB3 in a rapid spontaneous reaction. The hypothesis is that kinetochores (KT) further lower the activation energy. **C)** Analysis of mitotic checkpoint functionality in cells depleted of the Ndc80C or the Knl1C. HeLa cell were arrested in the G2 phase of the cell cycle, released into mitosis in the presence of 33 nM Nocodazole, and imaged six hours after release. The percentage of mitotic cells is shown. Collectively, 678, 270, and 588 cells were analyzed for the control, Ndc80C RNAi, and Knl1C RNAi conditions, respectively, in three repeats of the experiment. The red line represents the median. Scale bar = 10 µm.

While very slow *in vitro*, this conversion occurs rapidly at kinetochores, indicating that it may be catalyzed (Figure 1B) (Dick and Gerlich, 2013). Work *in vitro* and *in vivo* has now demonstrated that a scaffold of additional SAC proteins at the kinetochore acts as a recruiting platform for O-MAD2 and CDC20 and catalyzes their combination (Faesen et al., 2017; Kulukian et al., 2009; Lara-Gonzalez et al., 2021a; Piano et al., 2021). This scaffold comprises the MAD1:C-MAD2 core complex, the BUB1:BUB3 complex, and the MPS1 kinase (Figure 1A). MAD1, a coiled-coil protein unrelated to CDC20 except for the presence of a MIM motif, holds C-MAD2 in a remarkably stable 2:2 tetramer, the MAD1:C-MAD2 core complex (Chen et al., 1999; Sironi et al., 2002). The C-MAD2 molecules in the MAD1:MAD2 core complex recruit O-MAD2 from the cytosol through an asymmetric “conformational dimerization” (De Antoni et al., 2005; Mapelli et al., 2007). BUB1:BUB3 interacts with MAD1:C-MAD2, and together these proteins further recruit CDC20 to allow its rapid combination with O-MAD2 (Brady and Hardwick, 2000; Chen et al., 2023; Faesen et al., 2017; Fischer et al., 2021; Fischer et al., 2022; London and Biggins, 2014; Piano et al., 2021; Qian et al., 2017; Zhang et al., 2017). Swiveling of the C-terminal region of MAD1 is required for this conversion (Chen et al., 2023; Fischer et al., 2022; Heinrich et al., 2014; Kruse et al., 2014). These processes are collectively orchestrated by MPS1 kinase, which phosphorylates BUB1 to promote its interaction with MAD1, activates MAD1 through phosphorylation of its C-terminal region, and, in metazoans, promotes assembly of a specialized structure, the kinetochore corona, that cements these interactions before being dissolved by microtubule binding (Faesen et al., 2017; Fischer et al., 2021; Hewitt et al., 2010; Ji et al., 2017; London and Biggins, 2014; Qian et al., 2017; Raisch et al., 2022; Rodriguez-Rodriguez et al., 2018; Zhang et al., 2017). A set of SAC catalysts comprising MAD1:MAD2, BUB1:BUB3, and MPS1 is sufficient to promote a very significant acceleration of MCC assembly in an *in vitro* reconstituted system even in the absence of kinetochores (Faesen et al., 2017; Piano et al., 2021).

A single unattached kinetochore supports enough MCC assembly to maintain a mitotic delay (Dick and Gerlich, 2013; Rieder et al., 1995). Thus, we suspect that the catalytic efficiency of MCC assembly at kinetochores may be significantly higher than the one measured *in vitro* with isolated components. How kinetochores contribute to SAC signaling, however, remains unclear. All SAC components are recruited to the outer kinetochore, a 10-subunit assembly consisting of the KNL1, MIS12, and NDC80 subcomplexes, herewith respectively designated as Knl1C, Mis12C, and Ndc80C (Lara-Gonzalez et al., 2021b; Musacchio and Desai, 2017). Specifically, MPS1 kinase becomes recruited through an interaction with Ndc80C, which also serves as a microtubule-attachment platform through the NDC80^HEC1^ subunit (Hiruma et al., 2015; Ji et al., 2015; Pleuger et al., 2023; Sarangapani et al., 2021). Instead, BUB1:BUB3 becomes recruited by so-called MELT motifs in the KNL1 protein, after their phosphorylation by MPS1 kinase and recognition by BUB3, a phospho-aminoacid adaptor (London et al., 2012; Primorac et al., 2013; Shepperd et al., 2012; Yamagishi et al., 2012). Precisely how MAD1:MAD2 becomes recruited to kinetochores has remained partly elusive, but the already mentioned interaction with BUB1 contributes to this process, together with a binding site in the corona (Allan et al., 2020; Fischer et al., 2021; Heinrich et al., 2014; Kim et al., 2012; London and Biggins, 2014; Qian et al., 2017; Silio et al., 2015; Zhang et al., 2015).

How kinetochores influence MCC assembly is less clear. Depletion of KMN subunits has resulted in a range of different responses, likely because the penetrance of the depletion strongly influences the outcome, with partial depletions being compatible with a strong mitotic arrest, and complete depletion causing checkpoint override (Kim and Yu, 2015; Mariani et al., 2012; Martin-Lluesma et al., 2002; McCleland et al., 2003; Polley et al., 2023). Collectively, however, the current evidence is consistent with the notion that kinetochores are essential for SAC signaling. In addition to providing binding sites for the SAC catalysts, kinetochores may be expected to promote their productive interactions by orienting them favorably for their catalytic function on the MCC subunits. Furthermore, kinetochores are expected to couple microtubule binding to SAC suppression, a process whose molecular details are only beginning to be uncovered (Fischer, 2023; McAinsh and Kops, 2023). Here, we make new inroads into the dissection of these complex questions by combining biochemical *in vitro* reconstitutions of the human SAC and kinetochores. We demonstrate that kinetochores accelerate MCC accumulation also at high catalyst concentration, but become indispensable for rapid accumulation of MCC at physiological catalyst concentration. Our observations are consistent with a model in which both the concentration of SAC catalysts and their co-orientation promote efficient catalytic accumulation of MCC.

## Results and Discussion

### Kinetochores are required for the SAC response

To assess the role of kinetochores in the SAC response, we depleted by RNAi the outer kinetochore complexes Ndc80C or Knl1C. We then monitored the accumulation of mitotic cells six hours after their release from a G2 arrest in the presence of the spindle poison nocodazole, a potent SAC activator, counting the fraction of cells that had remained in mitosis with an active SAC (Figure 1C). More than 40 % of cells in the control condition were mitotic, as revealed by condensation of their chromosomes. Cells depleted of the outer kinetochore complexes, on the other hand, had all exited mitosis prematurely and had re-entered interphase. Thus, both Knl1C and Ndc80C are necessary to sustain the SAC. The SAC-promoting function of unattached kinetochores likely reflects their ability to recruit SAC proteins and to orient them reciprocally for catalytic MCC production. We therefore set out to shed light on this mechanism through biochemical reconstitution, harnessing our previous reconstitutions of SAC signaling and kinetochore organization with recombinant proteins (Faesen et al., 2017; Piano et al., 2021; Walstein et al., 2021; Weir et al., 2016).

### Engineering an inducible functional SAC scaffold

A crucial challenge towards achieving a functional biochemical reconstitution of kinetochores in SAC signaling is that the reconstituted kinetochore particles ought to recruit all SAC proteins. This goal, however, has not yet been fully achieved *in vitro*. A first challenge concerns MPS1, an apical kinase in SAC pathway. MPS1 contributes to SAC signaling in multiple ways. For instance, it phosphorylates the Met-Glu-Leu-Thr (MELT) motifs on KNL1 to recruit the BUB1:BUB3 complex (London et al., 2012; Primorac et al., 2013; Shepperd et al., 2012; Yamagishi et al., 2012). It also phosphorylates, after CDK1 priming phosphorylation, Thr461 of BUB1, located in the conserved motif 1 (CM1, residues 458-476), to promote the interaction of BUB1 with a conserved Arg-Leu-Lys (RLK) motif in the C-terminal coiled-coil region of MAD1 (Fischer et al., 2021; Ji et al., 2017; London and Biggins, 2014; Qian et al., 2017; Zhang et al., 2017). Additionally, MPS1 phosphorylates Thr716 in the MAD1 C-terminal RWD domain to facilitate binding to the N-terminal region of CDC20 (Faesen et al., 2017; Fischer et al., 2022; Ji et al., 2017). How MPS1 is recruited to kinetochores, however, remains unclear. Previous studies suggested that MPS1 binds to the Ndc80C competitively with microtubules, but there is evidence against this idea (Hayward et al., 2022; Hiruma et al., 2015; Ji et al., 2015; Vazquez-Novelle and Petronczki, 2010). Recent work in *S. cerevisiae* identified an MPS1 binding site in a region of Ndc80C also implicated in the interaction with the Dam1 complex (Parnell et al., 2024; Pleuger et al., 2024; Zahm and Harrison, 2024). Whether this site is functionally conserved in humans, however, is not yet clear. The implications of this are further discussed below. As we have not yet been able to reconstitute a robust MPS1-kinetochore interaction *in vitro* (SG and AM, unpublished results), for our *in vitro* experiments we either by pre-phosphorylating catalysts and kinetochores with MPS1, but also added MPS1 directly to an MCC assembly assay.

A second challenge concerns the recruitment of the MAD1:MAD2 “template” complex. MAD1:MAD2 recruitment requires interactions with BUB1 (facilitated by MPS1 phosphorylation) and with the building block of the kinetochore corona, the ROD-Zwilch-ZW10 (RZZ) complex, and is further regulated by Cyclin B (Alfonso-Perez et al., 2019; Allan et al., 2020; Fischer et al., 2021; Jackman et al., 2020; Ji et al., 2017; Klebig et al., 2009; Kops et al., 2005; Qian et al., 2017; Zhang et al., 2017; Zhang et al., 2015). Yet, we have so far been unable to identify a minimal set of interactions leading to robust MAD1:MAD2 recruitment to reconstituted kinetochores *in vitro* (SS and AM, unpublished observations). Kinetochore localization of MAD1:MAD2 signals checkpoint activation and mitotic arrest (Chen et al., 1998; Kuhn and Dumont, 2019; Waters et al., 1998). Ectopic retention of MAD1:MAD2 on bioriented kinetochores through a fusion with an outer kinetochore subunit is sufficient to cause metaphase arrest (Ballister et al., 2014; Heinrich et al., 2014; Kuijt et al., 2014; Maldonado and Kapoor, 2011). To target recombinant MAD1:MAD2 to reconstituted kinetochores, we engineered a strategy based on inducible dimerization (Ballister et al., 2014; Belshaw et al., 1996). We fused the FKBP-Rapamycin Binding (FRB) domain to the NUF2 subunit of the Ndc80C (Ndc80C^FRB^), and the 12-kDa FK506 binding protein (FKBP12, or simply FKBP) to the N-terminus of a MAD1 construct starting at residue 330 and extending to the entire C-terminus (^FKBP^MAD1^330-C^) (Figure 2A). This segment of MAD1 has been previously shown to be sufficient for SAC signaling *in vitro* (Faesen et al., 2017). We then induced the interaction between Ndc80C^FRB^ and ^FKBP^MAD1^330-C^ through addition of the small-molecular dimerizer Rapamycin (Belshaw et al., 1996). Analytical size-exclusion chromatography (SEC) confirmed that Ndc80C^FRB^ and ^FKBP^MAD1^330-C^:C-MAD2 formed a stoichiometric complex in the presence of Rapamycin (Figure 2B).

**Figure 2.**
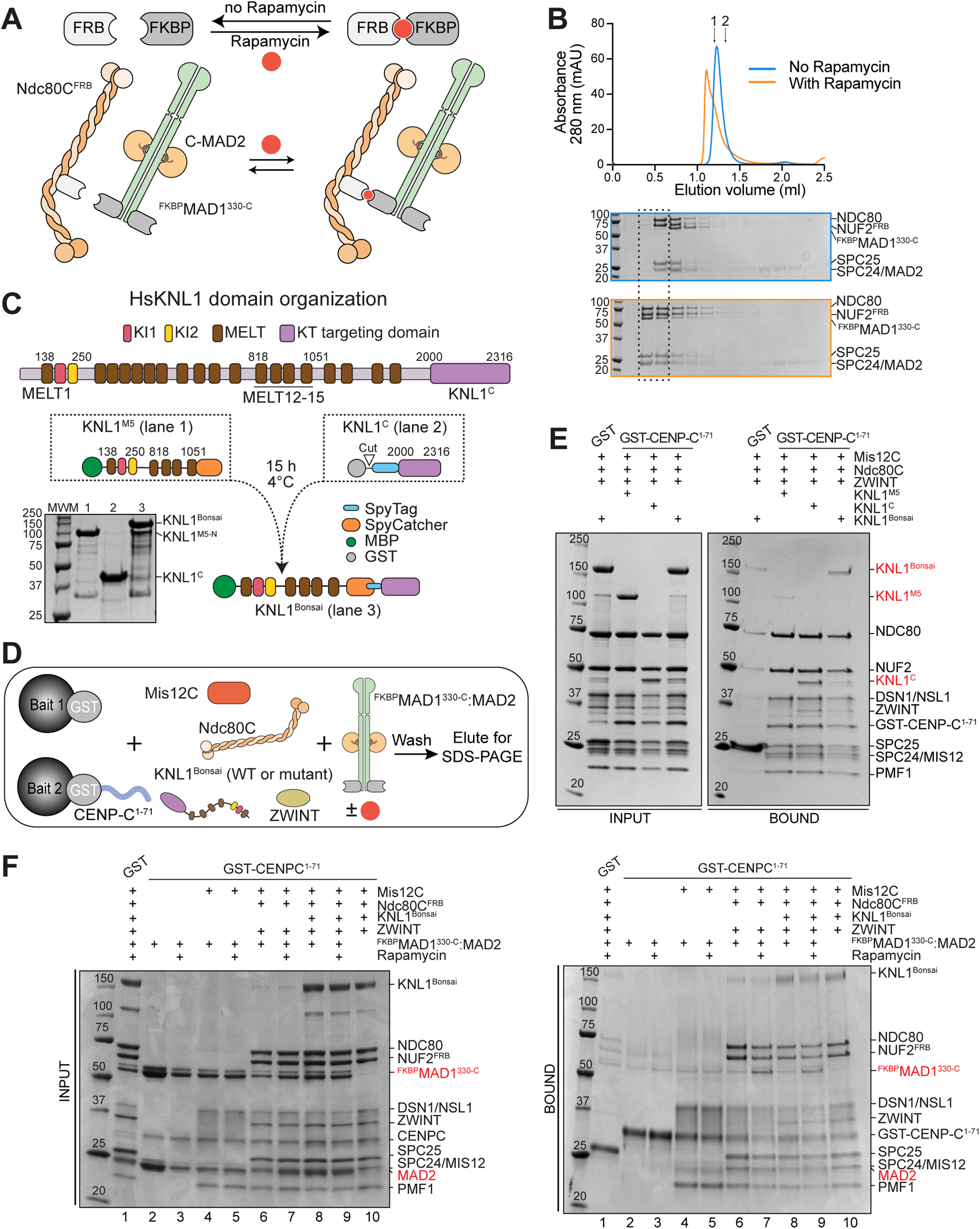
Engineering SAC-competent kinetochores in vitro. **A)** Schematic representation of FRB and FKBP dimerization through Rapamycin (red circle). In the presence of Rapamycin, of ^FKBP^MAD1^330-718^:MAD2 is expected to bind Ndc80C^FRB^, where the FRB moiety is fused at the C-terminus of the NUF2 subunit. **B)** Elution profiles and SDS PAGE analysis of fractions of size-exclusion chromatography experiments on a Superose 6 5/150 column with a stoichiometric mixture of ^FKBP^MAD1^330-718^:MAD2 and Ndc80C^FRB^ (3 μM) with or without 10 µM Rapamycin. Shown is a representative elution profile from three repeats. The arrow denoted “1” indicates the peak elution volume of the isolated Ndc80C, and the arrow denoted “2” indicates the peak elution volume of the isolated **C)** Scheme of motifs and domains of human KNL1. Numbering is according to isoform 2 of KNL1 (residues 1-2316, with residues 84-109 missing relative to the more canonical isoform 1 of 2342 residues). We opted to use isoform 2 numbering to facilitate a comparison with previously used constructs (Krenn et al., 2014). KI1 and KI2, lysine (K) – isoleucine (I) motifs; MELT, methionine (M) – glutamic acid (E) – leucine (L) – threonine (T) motifs. The KNL1^C^ fragment contains tandem RWD domains (RING finger-containing proteins, WD-repeat-containing proteins and yeast DEAD (DEXD)-like helicases (Petrovic et al., 2014). Strategy for generating KNL1^Bonsai^ through fusion of fragments fused to SpyTag or SpyCatcher. The KNL1^M5^ fragment is an engineered fusion of residues 138-250 and 818-1051. **D)** Scheme of solid phase binding assay monitoring interactions of outer kinetochore and SAC proteins. **E)** KNL1^C^ and KNL1^Bonsai^ bind Mis12C immobilized on solid phase, whereas only traces of KNL1^M5^ were detected, as expected. **F)** SDS PAGE result of pulldown with GST-CENP-C^1-71^ and ^R^KT with NUF2^FRB^ (3 μM) testing for binding of ^FKBP^MAD1:MAD2 (6 μM) with or without Rapamycin.

A third challenge concerns the recruitment of BUB1:BUB3. The BUB1:BUB3 complex binds through BUB3, a phospho-aminoacid adaptor, to so called MELT motifs on KNL1 after their phosphorylation by MPS1 (Figure 2C) (London et al., 2012; Primorac et al., 2013; Shepperd et al., 2012; Yamagishi et al., 2012). Our kinetochore reconstitutions so far included the C-terminal region of KNL1 (KNL1^C^), which binds the second Knl1C subunit ZWINT and mediates kinetochore targeting of the 2342-residue KNL1 protein (Kiyomitsu et al., 2007; Petrovic et al., 2014; Petrovic et al., 2010; Polley et al., 2024; Yatskevich et al., 2024). This region, however, lacks the MELT repeats (and KI motifs, see below), and is therefore unable to recruit BUB1:BUB3. To generate a KNL1 construct capable of recruiting BUB1:BUB3, we ligated two separately purified fragments of KNL1 encompassing the N- and C-terminal regions. One fragment, KNL1^M5^, embeds MELT1, the neighboring KI1 and KI2 motifs, and MELT repeats 12-15, which have been shown to be sufficient to sustain SAC signaling in cells (Vleugel et al., 2015).

The other segment, KNL1^C^, embeds the kinetochore-targeting domain (Figure 2C). We generated a mini-KNL1 protein (KNL1^Bonsai^) by covalent ligation of KNL1^M5^ and KNL1^C^ using a SpyTag/SpyCatcher pair (Zakeri et al., 2012) (Figure 2C). When electroporated into HeLa cells (Alex et al., 2019), a GFP-labelled version of KNL1^Bonsai^ (^GFP^KNL1^Bonsai^) decorated kinetochores in mitotic cells, indicating that our recombinant construct retains kinetochore localization properties similar to those of endogenous KNL1 (Figure S1A-C).

The N-terminal region of CENP-C provides a point of attachment for the KMN network to the kinetochore (Petrovic et al., 2016; Screpanti et al., 2011). To developed a binding assay to monitor the interaction of various moieties with the outer kinetochore, we immobilized a GST fusion of CENP-C^1-71^. We then monitored retention of outer kinetochore proteins on the solid phase (Figure 2D). GST-CENP-C^1-71^ bound the entire recombinant KMN network but failed to retain the KNL1^M5^ fragment. Conversely, both the KNL1^C^ and the KNL1^Bonsai^ constructs were effectively retained on solid phase (Figure 2E). Our reconstituted kinetochore incorporating KNL1^Bonsai^ (herewith indicated as ^R^KT, for “reconstituted kinetochore”) may also be expected to recruit BUB1:BUB3 upon phosphorylation of the five MELT repeats with MPS1. We demonstrate this prediction in the context of Figure 4. We also asked if ^R^KT, after immobilization on solid phase, recruited ^FKBP^MAD1^330-C^:C-MAD2 in the presence of Ndc80C^FRB^ and Rapamycin. Indeed, ^FKBP^MAD1^330-C^:C-MAD2 was retained on solid phase upon addition of Rapamycin (Figure 2F), indicating that we have obtained an inducible system for initiating SAC signaling on reconstituted kinetochores.

### Catalytic activation of SAC signaling by reconstituted kinetochores

Next, we tested if adding the ^R^KT to the SAC catalysts and the SAC substrates affected the rate of MCC production. For these experiments, we used our previously described FRET sensor consisting of BUBR1^mTurquoise^ as fluorescence donor and O-MAD2^TAMRA^ as fluorescence acceptor (Faesen et al., 2017). In this system, incorporation of MAD2 and BUBR1 in MCC is contingent on the presence of CDC20, so that the increase in FRET signal over time detects the time-dependent accumulation of MCC (Figure 3A).

**Figure 3.**
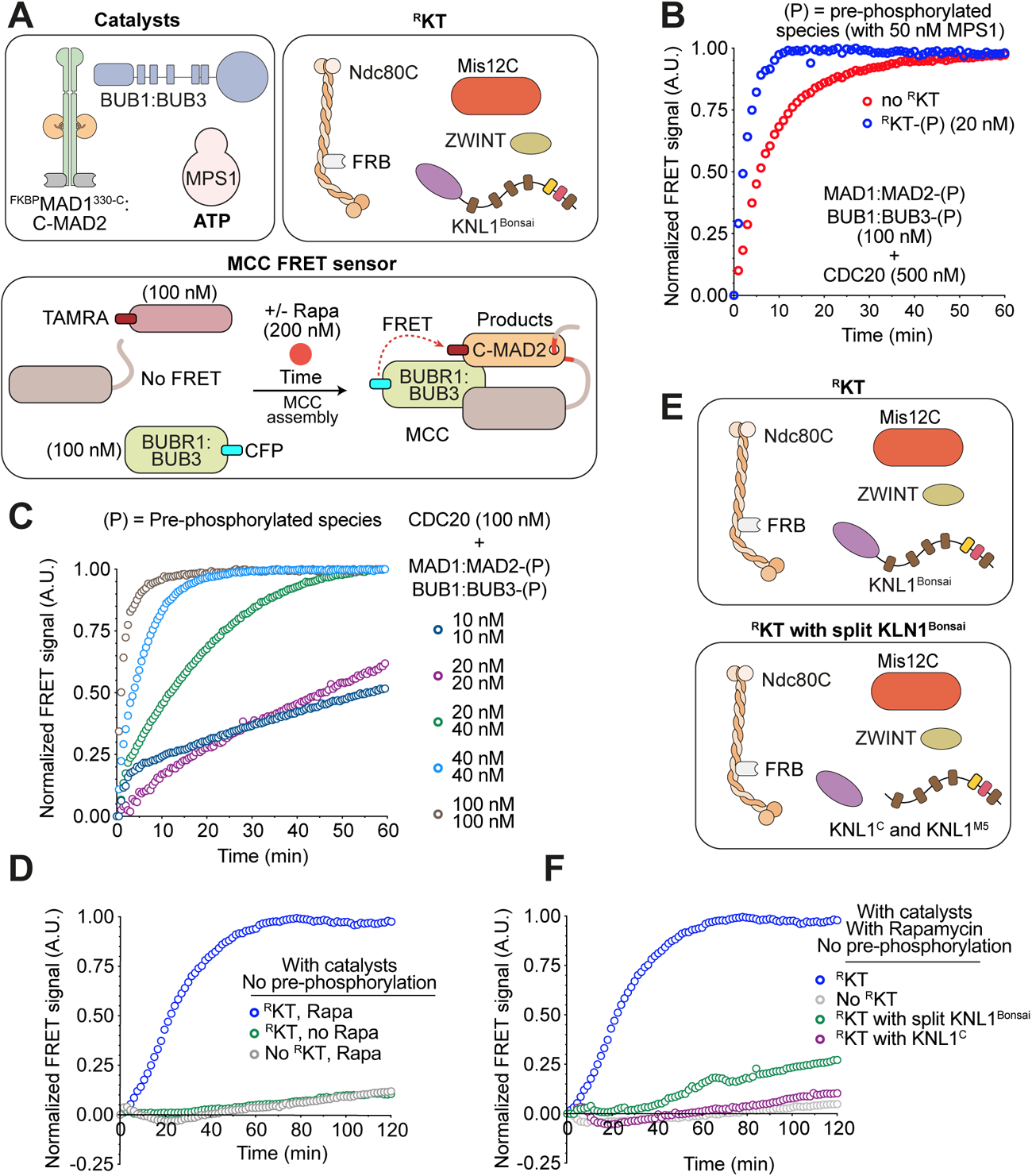
KMN supports catalytic MCC assembly. **A)** Scheme of catalysts, ^R^KT, and FRET sensor used to monitor MCC assembly (Faesen et al., 2017; Piano et al., 2021). ^CFP^BUBR1 and MAD2^TAMRA^ are donor and acceptor, respectively. CFP and TAMRA are closely positioned in MCC, allowing FRET (quantified as sensitized TAMRA fluorescence emission). **B)** Sensitized fluorescence emission (FRET) normalized to curve’s maximum for MCC sensor in the presence of the indicated concentration of MPS1-pre-phosphorylated catalysts, and supplemented with (blue) or without (red) pre-phosphorylated ^R^KTs (20 nM) in the presence of 200 nM Rapamycin. ATP (2 mM) and MgCl_2_ (10 mM) were added just before starting the experiment. All catalytic components were phosphorylated with MPS1 prior the assay. Unless otherwise specified, here and in other figures depicting FRET assays, the y-axis represents the fluorescence emission intensity of the FRET acceptor indicated as “Normalized FRET signal (A.U.),” where A.U. is arbitrary units. The blue and red curve were normalized to their own maximum value. **C)** Sensitized fluorescence emission (FRET) normalized to curve’s maximum for MCC sensor in the presence of the indicated concentration of MPS1-pre-phosphorylated catalysts. ^R^KTs were omitted. The dark blue and purple curves were not normalized as they did not reach a plateau during the observation time and their values were normalized to the maximum value of the light brown-color curve. **D)** Sensitized fluorescence emission (FRET) of MCC sensor in the presence of unphosphorylated catalysts (50 nM MPS1; 20 nM MAD1:MAD2; 40 nM BUB1:BUB3) and unphosphorylated ^R^KT (20 nM). The blue curve (also shown in panel **F**) was normalized to its own maximum. Fluorescence values of the other curves were normalized to the maximum value of the blue curve. The grey curve is also shown in Figure S1F. **E)** Variants of the ^R^KT used for controls in panel **F**. **F)** Sensitized fluorescence emission (FRET) of MCC sensor in the presence of unphosphorylated catalysts (same concentration as in panel **D**), unphosphorylated ^R^KT (same curve as in panel **D**), and the ^R^KT variants presented in panel **E**. Fluorescence values were normalized to maximum of the blue curve.

Previously, we demonstrated that addition of MAD1:MAD2 and BUB1:BUB3 (each at 100 nM) pre-phosphorylated with MPS1 kinase (50 nM) greatly accelerates MCC assembly in the presence of CDC20 (500 nM) (Faesen et al., 2017; Piano et al., 2021). Indeed, very rapid accumulation of MCC was observed under these conditions (Figure 3B, red curve. Table 1 reports concentrations of reagents used in MCC assembly assays throughout this work). Addition of ^R^KT (20 nM) that had also been pretreated with MPS1 (pre-phosphorylated ^R^KT, indicated as ^R^KT-(P)) to phosphorylate the five MELT motifs on KNL1^Bonsai^ further increased the rate of MCC production (Figure 3B, blue curve). The acceleration of the rate of MCC production by ^R^KT-(P)) was not complementary to that provided by catalysts, as it completely dependent on the presence of catalysts (Figure S1E). Thus, at an identical concentration of MCC substrates and pre-phosphorylated catalysts, addition of ^R^KT-(P) further accelerates MCC production, seemingly recapitulating *in vitro* a role of kinetochores in SAC signaling.

**Table 1.**
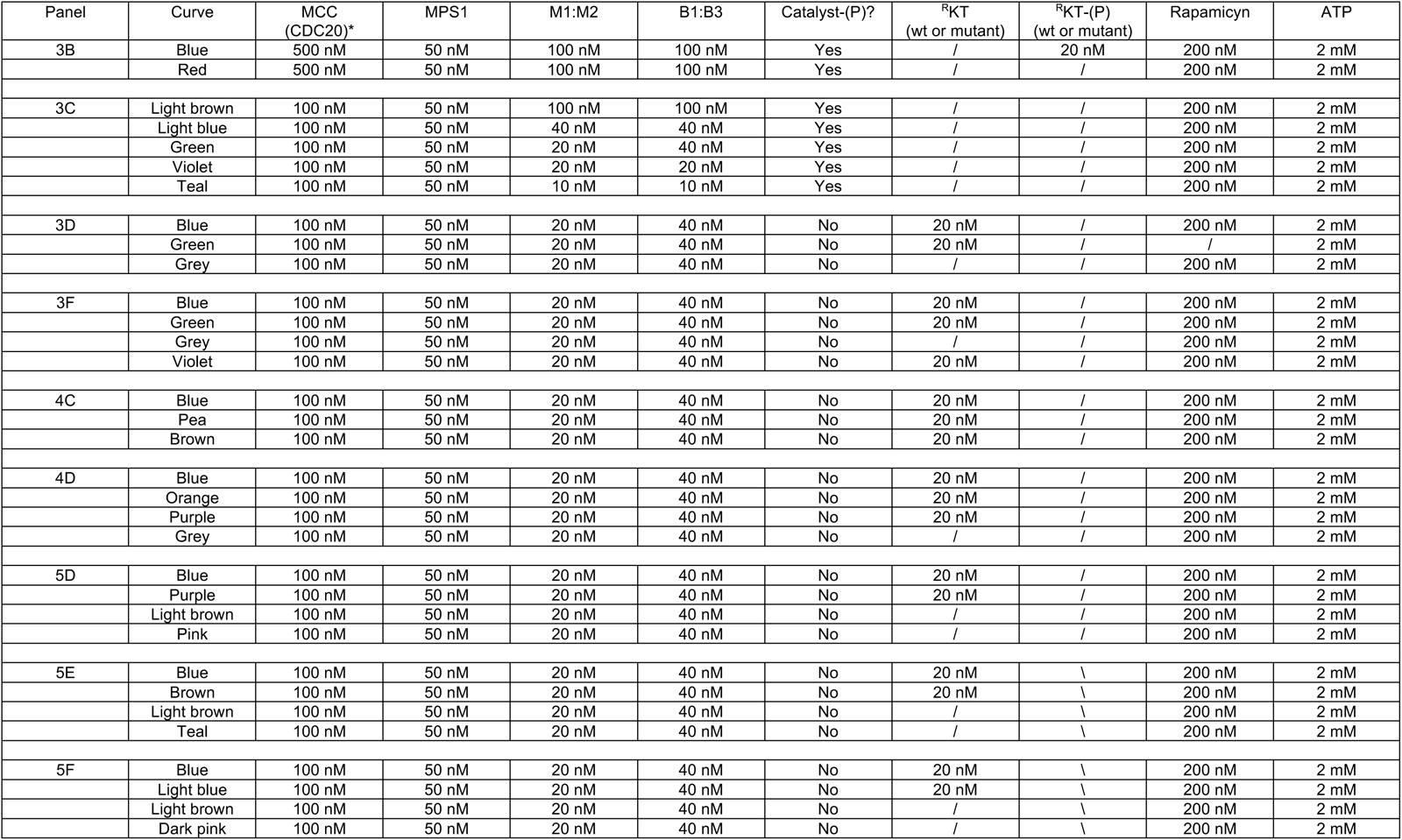
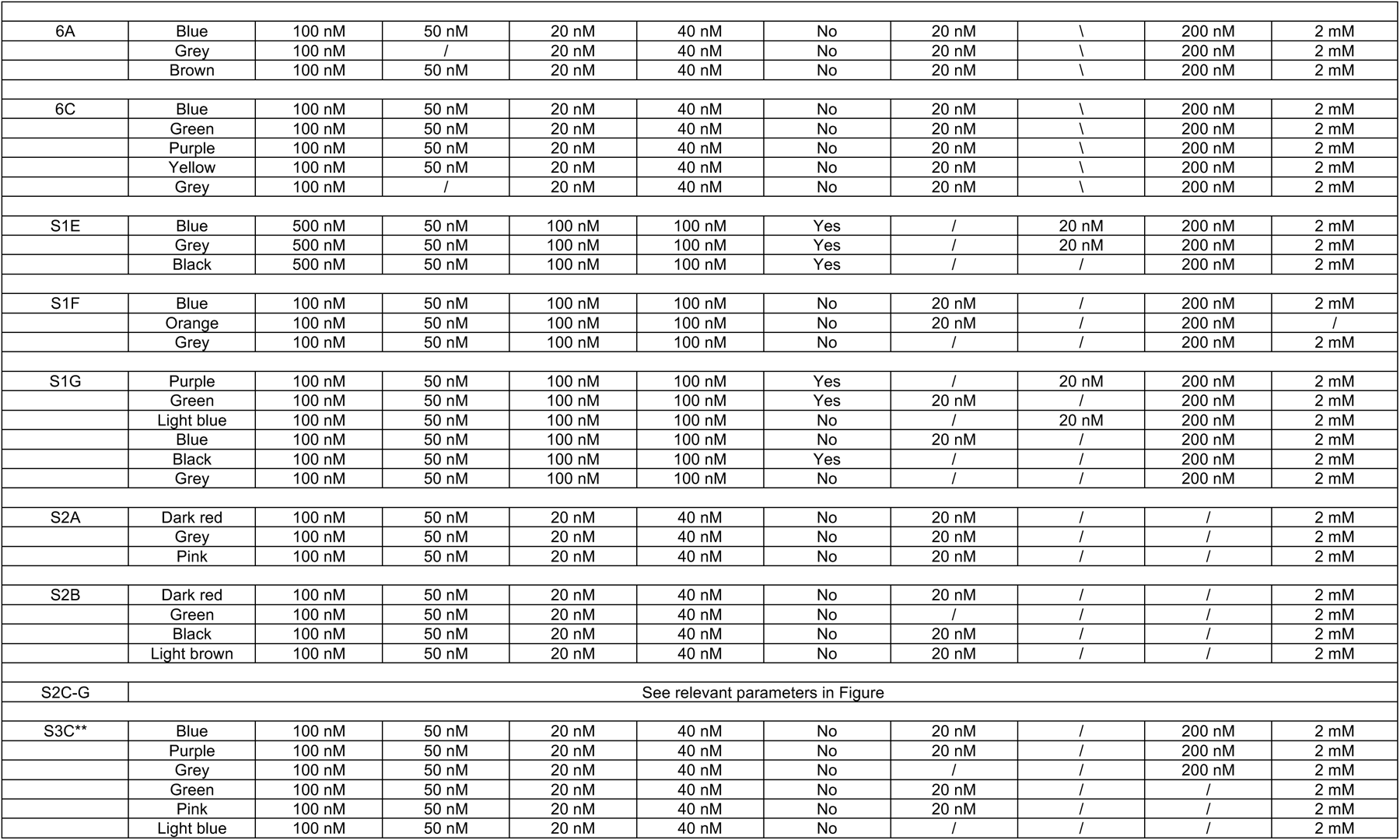
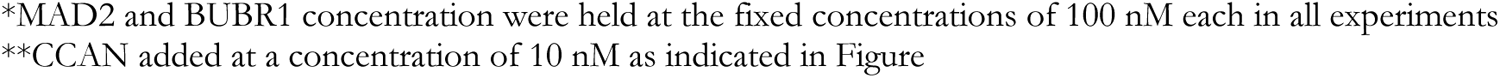
Concentration of species used in MCC assembly assays.

| Panel | Curve | MCC<br>(CDC20)* | MPS1 | M1:M2 | B1:B3 | Catalyst-(P)? | <sup>R</sup>KT<br>(wt or mutant) | <sup>R</sup>KT-(P)<br>(wt or mutant) | Rapamicyn | ATP |
| --- | --- | --- | --- | --- | --- | --- | --- | --- | --- | --- |
| 3B | Blue | 500 nM | 50 nM | 100 nM | 100 nM | Yes | / | 20 nM | 200 nM | 2 mM |
|  | Red | 500 nM | 50 nM | 100 nM | 100 nM | Yes | / | / | 200 nM | 2 mM |
| 3C | Light brown | 100 nM | 50 nM | 100 nM | 100 nM | Yes | / | / | 200 nM | 2 mM |
|  | Light blue | 100 nM | 50 nM | 40 nM | 40 nM | Yes | / | / | 200 nM | 2 mM |
|  | Green | 100 nM | 50 nM | 20 nM | 40 nM | Yes | / | / | 200 nM | 2 mM |
|  | Violet | 100 nM | 50 nM | 20 nM | 20 nM | Yes | / | / | 200 nM | 2 mM |
|  | Teal | 100 nM | 50 nM | 10 nM | 10 nM | Yes | / | / | 200 nM | 2 mM |
| 3D | Blue | 100 nM | 50 nM | 20 nM | 40 nM | No | 20 nM | / | 200 nM | 2 mM |
|  | Green | 100 nM | 50 nM | 20 nM | 40 nM | No | 20 nM | / | / | 2 mM |
|  | Grey | 100 nM | 50 nM | 20 nM | 40 nM | No | / | / | 200 nM | 2 mM |
| 3F | Blue | 100 nM | 50 nM | 20 nM | 40 nM | No | 20 nM | / | 200 nM | 2 mM |
|  | Green | 100 nM | 50 nM | 20 nM | 40 nM | No | 20 nM | / | 200 nM | 2 mM |
|  | Grey | 100 nM | 50 nM | 20 nM | 40 nM | No | / | / | 200 nM | 2 mM |
|  | Violet | 100 nM | 50 nM | 20 nM | 40 nM | No | 20 nM | / | 200 nM | 2 mM |
| 4C | Blue | 100 nM | 50 nM | 20 nM | 40 nM | No | 20 nM | / | 200 nM | 2 mM |
|  | Pea | 100 nM | 50 nM | 20 nM | 40 nM | No | 20 nM | / | 200 nM | 2 mM |
|  | Brown | 100 nM | 50 nM | 20 nM | 40 nM | No | 20 nM | / | 200 nM | 2 mM |
| 4D | Blue | 100 nM | 50 nM | 20 nM | 40 nM | No | 20 nM | / | 200 nM | 2 mM |
|  | Orange | 100 nM | 50 nM | 20 nM | 40 nM | No | 20 nM | / | 200 nM | 2 mM |
|  | Purple | 100 nM | 50 nM | 20 nM | 40 nM | No | 20 nM | / | 200 nM | 2 mM |
|  | Grey | 100 nM | 50 nM | 20 nM | 40 nM | No | / | / | 200 nM | 2 mM |
| 5D | Blue | 100 nM | 50 nM | 20 nM | 40 nM | No | 20 nM | / | 200 nM | 2 mM |
|  | Purple | 100 nM | 50 nM | 20 nM | 40 nM | No | 20 nM | / | 200 nM | 2 mM |
|  | Light brown | 100 nM | 50 nM | 20 nM | 40 nM | No | / | / | 200 nM | 2 mM |
|  | Pink | 100 nM | 50 nM | 20 nM | 40 nM | No | / | / | 200 nM | 2 mM |
| 5E | Blue | 100 nM | 50 nM | 20 nM | 40 nM | No | 20 nM | \ | 200 nM | 2 mM |
|  | Brown | 100 nM | 50 nM | 20 nM | 40 nM | No | 20 nM | \ | 200 nM | 2 mM |
|  | Light brown | 100 nM | 50 nM | 20 nM | 40 nM | No | / | \ | 200 nM | 2 mM |
|  | Teal | 100 nM | 50 nM | 20 nM | 40 nM | No | / | \ | 200 nM | 2 mM |
| 5F | Blue | 100 nM | 50 nM | 20 nM | 40 nM | No | 20 nM | \ | 200 nM | 2 mM |
|  | Light blue | 100 nM | 50 nM | 20 nM | 40 nM | No | 20 nM | \ | 200 nM | 2 mM |
|  | Light brown | 100 nM | 50 nM | 20 nM | 40 nM | No | / | \ | 200 nM | 2 mM |
|  | Dark pink | 100 nM | 50 nM | 20 nM | 40 nM | No | / | \ | 200 nM | 2 mM |
| 6A | Blue | 100 nM | 50 nM | 20 nM | 40 nM | No | 20 nM | \ | 200 nM | 2 mM |
|  | Grey | 100 nM | / | 20 nM | 40 nM | No | 20 nM | \ | 200 nM | 2 mM |
|  | Brown | 100 nM | 50 nM | 20 nM | 40 nM | No | 20 nM | \ | 200 nM | 2 mM |
| 6C | Blue | 100 nM | 50 nM | 20 nM | 40 nM | No | 20 nM | \ | 200 nM | 2 mM |
|  | Green | 100 nM | 50 nM | 20 nM | 40 nM | No | 20 nM | \ | 200 nM | 2 mM |
|  | Purple | 100 nM | 50 nM | 20 nM | 40 nM | No | 20 nM | \ | 200 nM | 2 mM |
|  | Yellow | 100 nM | 50 nM | 20 nM | 40 nM | No | 20 nM | \ | 200 nM | 2 mM |
|  | Grey | 100 nM | / | 20 nM | 40 nM | No | 20 nM | \ | 200 nM | 2 mM |
| S1E | Blue | 500 nM | 50 nM | 100 nM | 100 nM | Yes | / | 20 nM | 200 nM | 2 mM |
|  | Grey | 500 nM | 50 nM | 100 nM | 100 nM | Yes | / | 20 nM | 200 nM | 2 mM |
|  | Black | 500 nM | 50 nM | 100 nM | 100 nM | Yes | / | / | 200 nM | 2 mM |
| S1F | Blue | 100 nM | 50 nM | 100 nM | 100 nM | No | 20 nM | / | 200 nM | 2 mM |
|  | Orange | 100 nM | 50 nM | 100 nM | 100 nM | No | 20 nM | / | 200 nM | / |
|  | Grey | 100 nM | 50 nM | 100 nM | 100 nM | No | / | / | 200 nM | 2 mM |
| S1G | Purple | 100 nM | 50 nM | 100 nM | 100 nM | Yes | / | 20 nM | 200 nM | 2 mM |
|  | Green | 100 nM | 50 nM | 100 nM | 100 nM | Yes | 20 nM | / | 200 nM | 2 mM |
|  | Light blue | 100 nM | 50 nM | 100 nM | 100 nM | No | / | 20 nM | 200 nM | 2 mM |
|  | Blue | 100 nM | 50 nM | 100 nM | 100 nM | No | 20 nM | / | 200 nM | 2 mM |
|  | Black | 100 nM | 50 nM | 100 nM | 100 nM | Yes | / | / | 200 nM | 2 mM |
|  | Grey | 100 nM | 50 nM | 100 nM | 100 nM | No | / | / | 200 nM | 2 mM |
| S2A | Dark red | 100 nM | 50 nM | 20 nM | 40 nM | No | 20 nM | / | / | 2 mM |
|  | Grey | 100 nM | 50 nM | 20 nM | 40 nM | No | 20 nM | / | / | 2 mM |
|  | Pink | 100 nM | 50 nM | 20 nM | 40 nM | No | 20 nM | / | / | 2 mM |
| S2B | Dark red | 100 nM | 50 nM | 20 nM | 40 nM | No | 20 nM | / | / | 2 mM |
|  | Green | 100 nM | 50 nM | 20 nM | 40 nM | No | / | / | / | 2 mM |
|  | Black | 100 nM | 50 nM | 20 nM | 40 nM | No | 20 nM | / | / | 2 mM |
|  | Light brown | 100 nM | 50 nM | 20 nM | 40 nM | No | 20 nM | / | / | 2 mM |
| S2C-G | See relevant parameters in Figure |  |  |  |  |  |  |  |  |  |
| S3C** | Blue | 100 nM | 50 nM | 20 nM | 40 nM | No | 20 nM | / | 200 nM | 2 mM |
|  | Purple | 100 nM | 50 nM | 20 nM | 40 nM | No | 20 nM | / | 200 nM | 2 mM |
|  | Grey | 100 nM | 50 nM | 20 nM | 40 nM | No | / | / | 200 nM | 2 mM |
|  | Green | 100 nM | 50 nM | 20 nM | 40 nM | No | 20 nM | / | / | 2 mM |
|  | Pink | 100 nM | 50 nM | 20 nM | 40 nM | No | 20 nM | / | / | 2 mM |
|  | Light blue | 100 nM | 50 nM | 20 nM | 40 nM | No | / | / | / | 2 mM |

\*MAD2 and BUBR1 concentration were held at the fixed concentrations of 100 nM each in all experiments
\*\*CCAN added at a concentration of 10 nM as indicated in Figure

Under the conditions that support rapid accumulation of MCC even in the absence of kinetochores in our FRET assay ((Faesen et al., 2017; Piano et al., 2021) and Figure 3B), the concentrations of catalysts are higher than those estimated for human cells. MAD1, for instance, may be present at 10-20 nM in cells (Luo et al., 2004; Shah et al., 2004) but was used at 100 nM in Figure 3B. To assess how the rate of MCC accumulation scales with catalyst concentration, we pre-phosphorylated the catalysts with MPS1 and diluted them in the MCC FRET assays. MCC accumulation progressed very rapidly at the highest concentrations of catalysts (40 nM or 100 nM for both MAD1:MAD2 and BUB1:BUB3), apparently insensitive to the decrease of CDC20 concentration from 500 nM to 100 nM (compare experiments in Figure 3B-C), i.e. a concentration equimolar with the concentration of MAD2 and BUBR1:BUB3 and closer to the cellular concentration of CDC20, estimated between 100 and 285 nM (Fang, 2002; Tang et al., 2001). On the other hand, the rate of MCC assembly decreased considerably at lower concentrations of pre-phosphorylated catalysts (Figure 3C).

At 20 nM MAD1:MAD2 and 40 nM BUB1:BUB3, an intermediate rate of stimulation of MCC assembly was observed (Figure 3C). We settled on this concentration of catalysts to test the role of pre-phosphorylation and kinetochores in MCC assembly. We reasoned that pre-phosphorylation may bypass a requirement for kinetochore-localized kinase activity for effective accumulation of phosphorylated scaffolds and catalysts. To test this, we monitored the rate of MCC production in the absence of MPS1 pre-phosphorylation of catalysts. In the absence of pre-phosphorylation, a concentration of catalysts of 20 nM MAD1:MAD2 and 40 nM BUB1:BUB3 failed to promote rapid accumulation of MCC assembly, probably because MPS1-mediated phosphorylation of these reagents at these concentration proceeds too slowly (Figure 3D, grey curve). Remarkably, addition of 20 nM ^R^KT strongly stimulated MCC assembly (Figure 3D, blue curve). Stimulation was dependent on addition of Rapamycin, indicating that it requires MAD1:MAD2 accumulation on kinetochores (green curve in Figure 3D and Figure S2A-B). It was also dependent on ATP, required for kinase activity (Figure S1F). Stimulation also required intact KNL1^Bonsai^, because ^R^KTs incorporating unfused KNL1^C^ and KNL1^M5^ (split KNL1^Bonsai^) did not support accelerated MCC production (Figure 3F; green curve). Finally, K^C^MN was unable to stimulate MCC assembly, indicating that functional MELT repeats on the K^M5^ fragment are required (Figure 3F, purple curve).

Collectively, these observations are consistent with the idea that incorporation in the ^R^KT of MAD1:MAD2 (through Rapamycin-induced binding to Ndc80C) and BUB1:BUB3 (through MPS1-mediated phosphorylation of MELT repeats) greatly enhances the rate of MCC assembly. Confirming the importance of MELT phosphorylation by MPS1, adding ^R^KT-(P) to catalysts at the standard concentration resulted in much more rapid MCC production in comparison to when ^R^KT was not pre-phosphorylated (Figure S1G). Even if not pre-phosphorylated, however, the ^R^KT accelerated MCC assembly relative to a condition in which it was omitted altogether (Figure S1G). Omission of the ^R^KT in the absence of pre-phosphorylation of the catalysts did not allow rapid assembly of MCC (Figure S1G), even at very high concentrations of catalysts (Figure S2C-G).

The constitutive centromere-associated network (CCAN) is a group of 16 proteins in humans that connects the outer kinetochore KMN network to the centromere (Yatskevich et al., 2023). We modified our solid phase binding assay by replacing CENP-C^1-71^ with CENP-C^1-544^ (Figure S3A). In addition to binding Mis12C, this segment of CENP-C also binds the CCAN complex (Hinshaw and Harrison, 2013; Klare et al., 2015; McKinley et al., 2015; Nagpal et al., 2015; Pesenti et al., 2022; Tian et al., 2022; Yatskevich et al., 2022). As expected, both the ^R^KT and CENP11, a CCAN sub-assembly consisting of 11 subunits previously shown to be sufficient to bind CENP-C (the 12^th^ CCAN subunit) (Pesenti et al., 2018), were retained on solid phase in the presence of CENP-C^1-544^ (Figure S3B). To assess a possible role of CCAN in MCC assembly, we ran the MCC assembly assay in the presence of catalysts, ^R^KT, and Rapamycin and asked whether addition of CENP12 (including CENP-C^1-544^) enhanced the MCC assembly rate. There was no difference in the rate of assembly of MCC with or without CENP12, suggesting that the latter does not stimulate MCC assembly (Figure S3C).

### Relative potency of MELT and KI motifs of KNL1

The MELT repeat unit is flanked by additional motifs, including the Thr-X-X-Ω (X = any aminoacid, Ω = Phe or Tyr) and Ser-His-Thr (SHT) motifs, positioned N- and C-terminally to the MELT sequence, respectively. The SHT motif contributes to recruiting BUB1:BUB3 to the kinetochore (Vleugel et al., 2015). Two additional motifs, KI1 and KI2, interact respectively with the tetratricopeptide repeat (TPR) domains of BUB1 and BUBR1. Together with the first MELT motif (MELT1) of KNL1, the KI1 and KI2 motifs form a reinforced unit for BUB1:BUB3 recruitment and SAC signaling (Bolanos-Garcia et al., 2011; Kiyomitsu et al., 2007; Krenn et al., 2014; Krenn et al., 2012; Vleugel et al., 2013). We carried out additional experiments to assess the role of MELT phosphorylation in the catalytic assembly of MCC. First, we generated a set of KNL1 variants and mutants as indicated in Figure 4A, mixed them with Mis12C and Ndc80C, and immobilized them on solid phase through GST-CENP-C^1-71^. Then, we tested the ability of the various KNL1 species to bind and retain BUB1:BUB3 on solid phase upon phosphorylation with MPS1 kinase. KNL1^C^ did not bind BUB1:BUB3 above background levels, as expected for a construct lacking MELT motifs altogether (Figure 4B, lane 2; the negative control is in lane 1). Conversely, KNL1^Bonsai^ or its MPS1 pre-phosphorylated form bound BUB1:BUB3, with pre-phosphorylation leading to apparently superstoichiometric binding (lanes 3-4). Without MPS1 pre-phosphorylation, mutation of the KI1 and KI2 motifs (K^KI1+2^MN) led to a strong decrease of BUB1:BUB3 binding (lane 5), consistent with the idea that KI1 and KI2 provide a high-affinity, phosphorylation-independent binding site for BUB1:BUB3 (Kiyomitsu et al., 2007; Krenn et al., 2014; Krenn et al., 2012). MPS1 pre-phosphorylation of K^KI1+2^MN enhanced BUB1:BUB3 binding, likely through phosphorylation of the MELT motifs (lane 6). Finally, a mutant in which both the MELT and the adjacent SHT motifs were mutated (to MELA and AHA, respectively; K^M5-A^MN) was also a weak BUB1:BUB3 binder, and as expected pre-phosphorylation did not rescue this effect of the mutations, confirming that MPS1 phosphorylation promotes BUB1:BUB3 binding through the MELT repeats (lanes 7-8). Collectively, these binding experiment indicate that ^R^KTs containing KNL1^Bonsai^ and its variants behave as expected based on previous binding analyses (Jema et al., 2023; Ji et al., 2017; Kiyomitsu et al., 2007; Krenn et al., 2014; London et al., 2012; Roy et al., 2020; Shepperd et al., 2012; Vleugel et al., 2015; Vleugel et al., 2013; Yamagishi et al., 2012; Zhang et al., 2014).

Next, we assessed how different variants of KNL1^Bonsai^ in ^R^KTs affected the rate of MCC accumulation in our FRET assay. Both the KNL1^M5/A^ and the KNL1^KI1+2/A^ mutants caused a strong reduction of the rate of MCC accumulation (Figure 4C; the residual activation was dependent on Rapamycin, as shown in Figure S2A-B). These results imply that the KI1 and KI2 motifs contribute substantially to catalytic MCC assembly in the context of KNL1^Bonsai^, and that they play a role in MCC assembly even in the presence of five MELT motifs in this construct. The results are consistent with the idea that the combination of closely spaced KI1 and MELT1 creates a composite binding site that binds BUB1:BUB3 cooperatively and with high affinity, decreasing its turnover and providing a stable platform for the SAC catalytic apparatus, as previously suggested by an *in vivo* analysis (Krenn et al., 2014). In agreement with this idea, a construct containing only the first MELT repeat and the two adjacent KI motifs (KNL1^Bonsai-M1^, generated as indicated schematically in Figure S4A), bound BUB1 on solid phase (Figure S4B-C) and was a robust activator of MCC assembly in the presence of Rapamycin but not in its absence (Figure 4D and Figure S2A-B).

**Figure 4.**
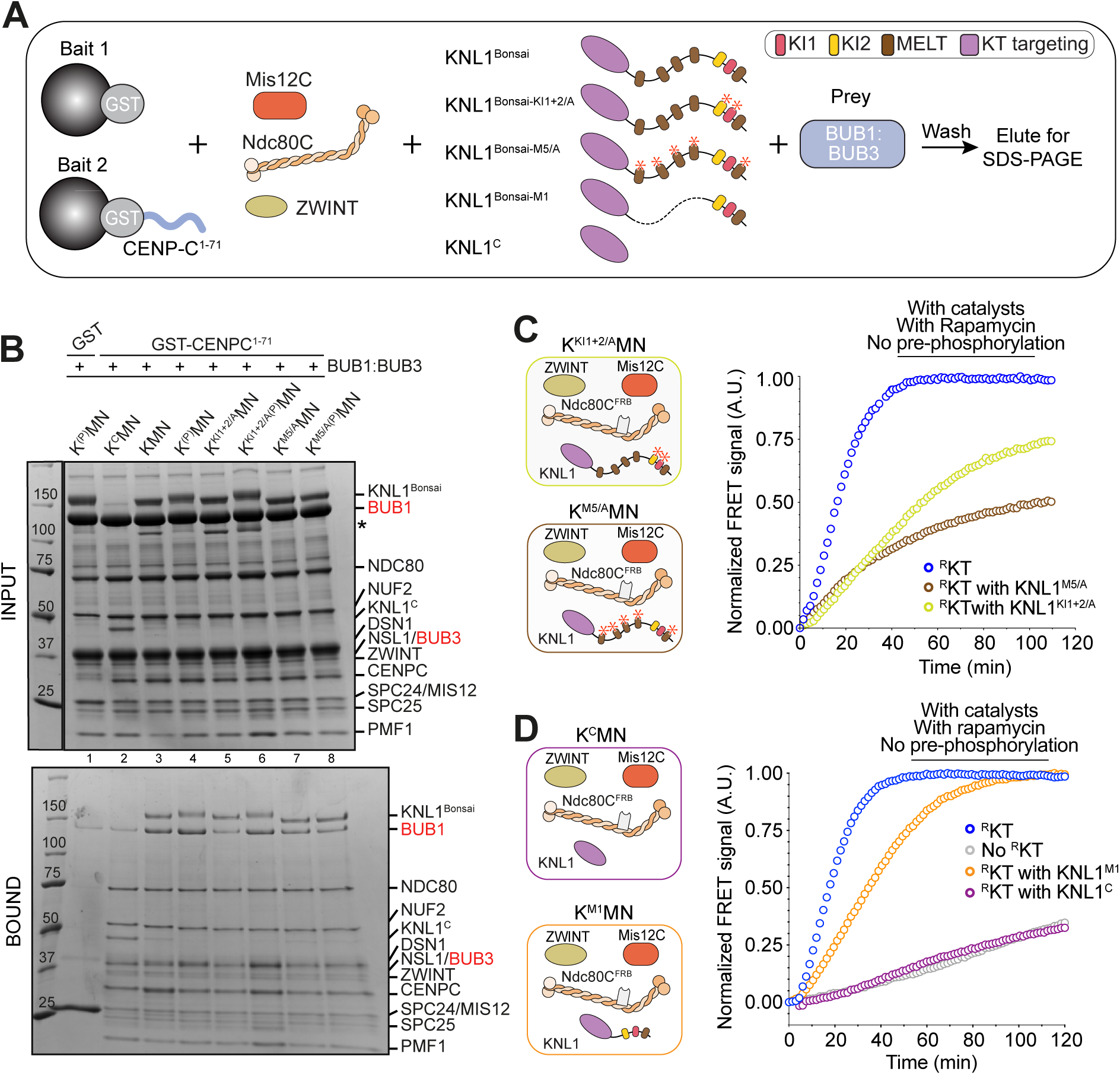
KNL1^Bonsai^ brings BUB1:BUB3 in proximity of MAD1:MAD2. **A)** Scheme of GST pulldown assays with CENP-C^1-71^ and ^R^KT (3 µM) assembled with the indicated versions of Knl1C and further incubated with BUB1:BUB3 (10 µM) as pray. **B)** SDS PAGE of pulldown experiment schematized in panel **A** and containing GST-CENP-C^1-71^ and ^R^KT at 3 µM with different KNL1 mutants and BUB1:BUB3 at 10 µM as indicated above each lane. The asterisks indicate excess of KNL1^M5^ fragments not fused to KNL1^C^. **C)** Sensitized fluorescence emission (FRET) of MCC sensor in the presence of unphosphorylated catalysts and unphosphorylated ^R^KT containing the indicated KNL1 variants, supplemented at the same concentrations used in Figure 3D. The blue curve was normalized to its maximum. The other curves did not plateau during the experiment and were normalized to the maximum of the blue curve. **D)** Sensitized fluorescence emission (FRET) of MCC sensor in the presence of unphosphorylated catalysts and unphosphorylated ^R^KT containing the additional indicated KNL1 variants, supplemented at the same concentrations used in Figure 3D. The blue and orange curves were normalized to their own maximum. The grey and purple curves were normalized to the maximum of the blue curve.

### Role of BUB1 in MCC assembly

BUB1 contains sequence motifs that promote interactions with the kinetochore and with other SAC components (Figure S4D), and that contribute to catalytic MCC assembly in the absence of reconstituted kinetochores (Piano et al., 2021). We were curious to verify if these motifs become dispensable in the presence of kinetochores. Interaction of two single-helices promotes the interaction of the BUB1 and BUBR1 paralogs and kinetochore recruitment of BUBR1 (Overlack et al., 2015). In our GST-CENP-C^1-71^ pulldown assay, a mutant lacking the helix (BUB1^Δhelix^) failed to recruit BUBR1 (Figure 5A-B). Replacing BUB1 with BUB1^Δhelix^ did not cause any effect on the rate of MCC formation in the presence of ^R^KT *in vitro* (Figure 5C-D). This result recapitulates our previous results *in vivo* (Overlack et al., 2017; Overlack et al., 2015) and demonstrates that the recruitment of BUBR1:BUB3 to the catalytic platform is not required for rapid MCC assembly.

**Figure 5.**
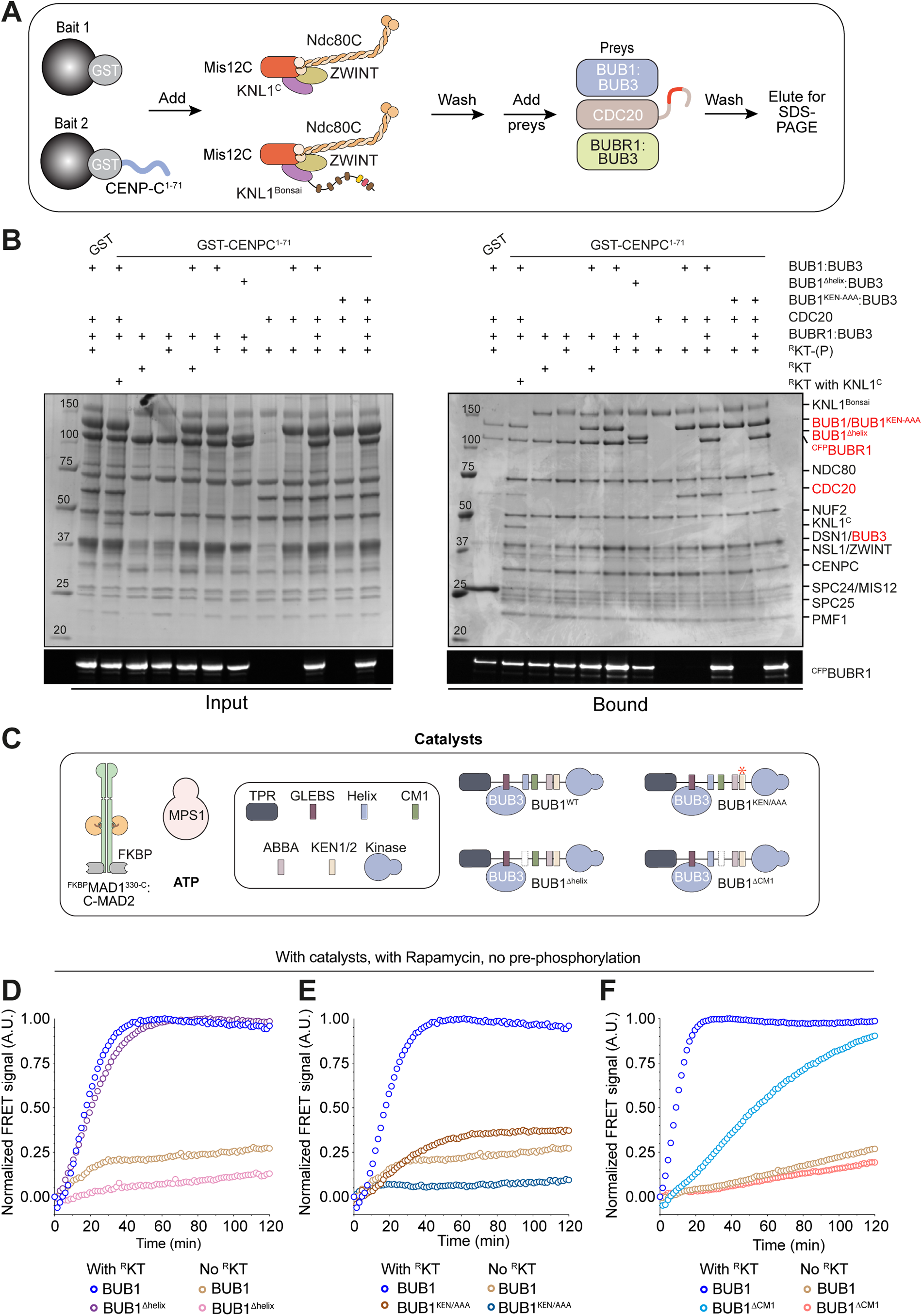
Influence of BUB1:BUB3 interaction with other SAC components in catalysis. **A)** Scheme of pulldown assays with CENP-C^1-71^, ^R^KT or ^R^KT assembled with KNL1^C^ (rather than KNL1^Bonsai^, each at 3 μM), and prays including BUB1:BUB3 (10 μM), BUBR1:BUB3 (8 μM), and CDC20 (6 μM). **B)** SDS PAGE of pulldowns with baits and preys indicated above each lane. Pre-phosphorylation of the ^R^KTs species by MPS1 (50 nM) is indicated with (P). Coomassie staining is shown above in-gel fluorescence (488 nm) below. All baits were incubated with an at least 3-fold molar excess of BUB1:BUB3, BUBR1:BUB3, and CDC20. Efficient phosphorylation of KNL1 and KNL1^KI1+2/AA^ was confirmed by a shift in migration in SDS-PAGE of the KNL1 band. No shift of KNL1^M5-A^ upon phosphorylation with MPS1 was observed. **C)** Scheme of the experimental setup of the following assays. **D)** MCC FRET assay monitoring the assembly of MCC without pre-phosphorylation of the components by MPS1, in the presence of the catalysts wild-type, Rapamycin with (blue curve, also shown in panel **E**) and without ^R^KT (light brown curve, also shown in panel **E**), and with BUB1^Δhelix^mutant with (thistle curve) and without ^R^KT (pink curve). **E)** MCC FRET assay monitoring the assembly of MCC without pre-phosphorylation of the components by MPS1, in the presence of the catalysts wild-type, Rapamycin with (blue curve, already shown in panel **D**) and without ^R^KT (light brown curve, already shown in panel **D**), and with BUB1^KEN/AAA^ mutant with (dark brown curve) and without ^R^KT (dark blue curve). **F)** MCC FRET assay monitoring the assembly of MCC without pre-phosphorylation of the components by MPS1, in the presence of the catalysts wild-type, Rapamycin with (blue curve) and without ^R^KT (light brown curve), and with BUB1^ΔCM1^ mutant with (light blue curve) and without ^R^KT (salmon curve).

Using its ABBA, KEN, and CM1 motifs, BUB1 positions CDC20 in close proximity to the MAD1 coiled-coil (Lara-Gonzalez et al., 2021a; Piano et al., 2021). In our pulldown assay, a BUB1^KEN/AAA^ mutant (where both the KEN and ABBA motifs were mutated to alanine) retained lower amounts of CDC20 relative to wild type BUB1. In the MCC assembly assay, the same mutant demonstrated a dramatic decrease in the rate of MCC assembly (Figure 5C,E). The CM1, a target of CDK1:Cyclin B and MPS1 phosphorylation, contributes to MAD1:MAD2 recruitment to the kinetochore (Fischer et al., 2021; Fischer et al., 2022; Ji et al., 2017; Klebig et al., 2009; London and Biggins, 2014; Qian et al., 2017; Zhang et al., 2017). In the MCC assembly assay, deletion of CM1 led to a significant reduction in the rate of MCC assembly, but did not abrogate catalytic MCC assembly (Figure 5C,F). *In vivo*, inhibition of the BUB1:MAD1 binding interaction is incompatible with robust SAC signaling (Ballister et al., 2014; Heinrich et al., 2014; London and Biggins, 2014; Silio et al., 2015). Nonetheless, our observation that MCC assembly is only partially impaired when MAD1 and BUB1 cannot interact suggests that kinetochores, by recruiting BUB1 and MAD1:MAD2 in close proximity, compensate for the loss of an interaction of the catalysts.

### BUB1 and MAD1:MAD2 are an integrated platform for catalytic MCC assembly by MPS1

As discussed earlier, phosphorylation of KNL1 by MPS1 is required for a robust rate of MCC assembly. MPS1 has also been shown to phosphorylate T461 in the CM1 motif of BUB1, which in turn promotes the interaction with MAD1 and its kinetochore recruitment (Fischer et al., 2021; Ji et al., 2017; London and Biggins, 2014; Qian et al., 2017). In addition, MPS1 phosphorylates residue T716 at the C-terminus of MAD1, an event that activates the catalytic function of MAD1 (Allan et al., 2020; Chen et al., 2023; Faesen et al., 2017; Ji et al., 2017; Piano et al., 2021). To investigate the specific role of MPS1 kinase activity in our MCC assembly assay, we conducted a comprehensive analysis to dissect individual phosphorylation requirements for this kinase. Addition of Reversine, a potent inhibitor of MPS1 kinase activity (Santaguida et al., 2010) ablated catalytic MCC assembly (Figure 6A). At high concentrations of pre-phosphorylated catalysts and in the absence of kinetochores, individual alanine mutations of Ser459 and Thr461 on BUB1 did not inhibit MCC catalysis *in vitro* (Piano et al., 2021). We combined the mutations in a single mutant and performed the MCC assembly reaction in the presence of kinetochores. The mutations, when combined, caused a moderate decline in the rate of MCC assembly (Figure 6B-C). Additionally, when we replaced wild-type MAD1 with the MAD1^T716A^ mutant in the MCC assembly assay, we also observed a pronounced decrease of the rate of MCC assembly (Figure 6B-C). Finally, when the phospho-alanine mutants MAD1^T716A^ and BUB1^S459A-T461A^ were combined, the rate of MCC assembly was similar to that in the absence of MPS1 kinase (Figure 6B-C).

**Figure 6.**
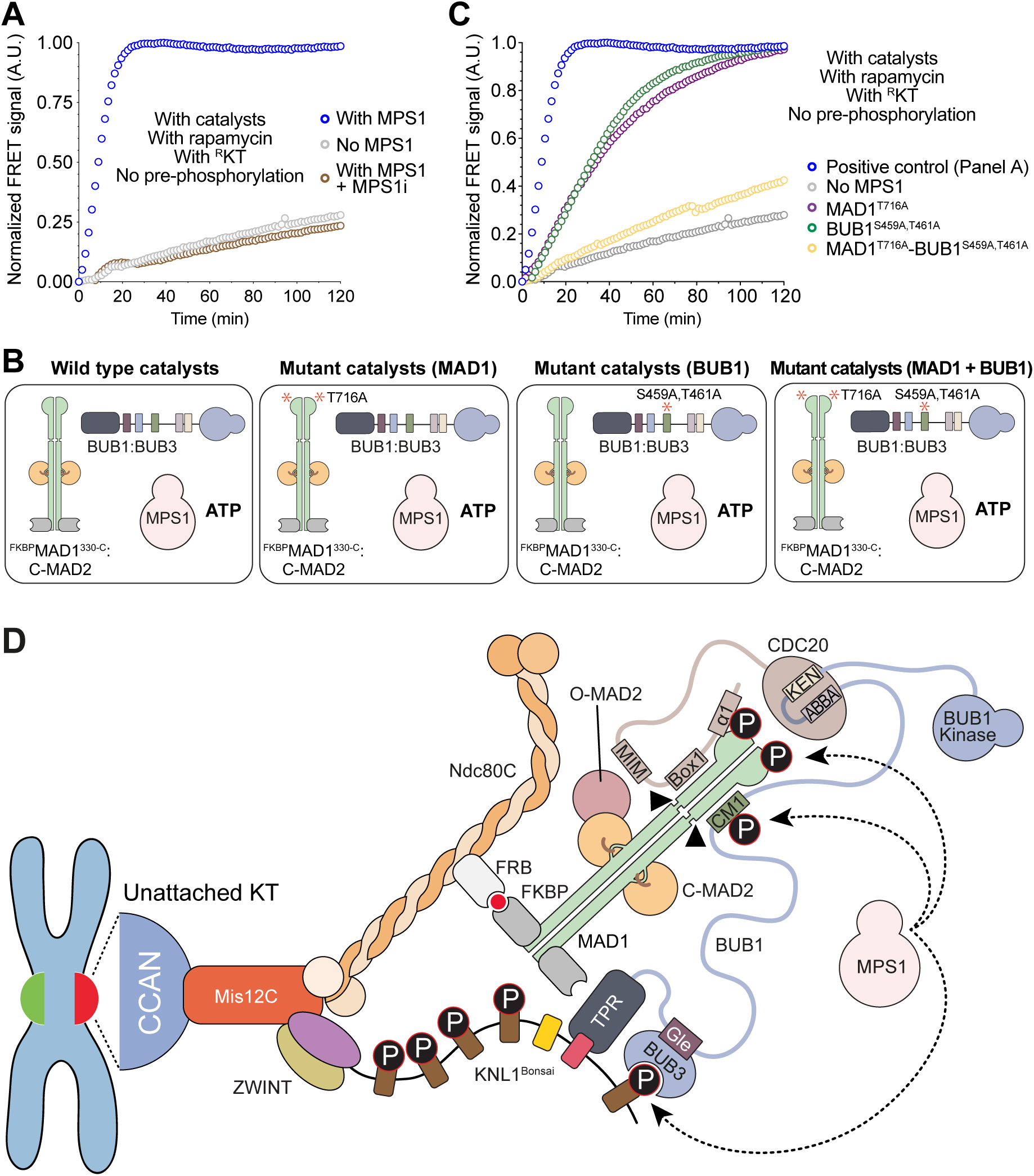
BUB1 and MAD1:MAD2 are an integrated platform for catalytic MCC assembly by MPS1. **A)** MCC FRET assay monitoring the assembly of MCC without pre-phosphorylation of the components by MPS1, in the presence of the catalysts wild-type, Rapamycin with ^R^KT (blue curve, already shown in Figure 5F), without MPS1 (grey curve, also shown in panel **C**) and in the presence of Reversine, MPS1 inhibitor (brown curve). **B)** Schematic representation of the experimental setup of the following assays. **C)** MCC FRET assay monitoring the assembly of MCC without pre-phosphorylation of the components by MPS1, in the presence of Rapamycin and ^R^KT (blue curve, already shown in Figure 5F), and with wild-type or mutant catalysts: BUB1^S459A-T461A^ (green curve), MAD1^T716A^ (purple curve), both BUB1^S459A-T461A^ and MAD1^T716A^ (yellow curve), or without MPS1 (grey curve, already shown in panel **A**). **D)** Schematic representation of the assembly of the catalytic scaffold on the engineered outer kinetochore, highlighting the crucial binding interfaces and phosphorylation sites for efficient MCC formation.

### Conclusions

In this study, we have pushed the envelope for *in vitro* biochemical reconstitutions by combining two major reconstitutions in a single test tube, a reconstitution of the kinetochore and a reconstitution of the SAC. This approach allowed us to investigate the effects of the kinetochore on SAC signaling. After solving several challenges to allow robust recruitment of SAC components to ^R^KT, we were able to demonstrate a role of ^R^KTs in the acceleration of MCC assembly. This acceleration is contingent on the recruitment of MAD1:MAD2 and BUB1:BUB3 to ^R^KTs, as it requires Rapamycin (to recruit MAD1:MAD2) and the MELT and KI repeats of KNL1 (to recruit BUB1:BUB3). Furthermore, acceleration of MCC assembly is contingent on the interaction of MAD1:MAD2 and BUB1:BUB3, as mutations in the CM1 motif of BUB1 affect the rate of MCC assembly. Thus, even within ^R^KT, the juxtaposition of MAD1:MAD2 and BUB1:BUB3 contributes substantially to catalysis (Figure 6D), indicating that kinetochores do not simply work by concentrating different SAC catalysts in a single scaffold.

We resorted to an artificial dimerization system to trigger MAD1:MAD2 recruitment to ^R^KTs, as we have not yet been able to recapitulate robust recruitment of MAD1:MAD2 with unmodified purified components. While the precise reasons for this will require further investigations, we suspect that introduction of a proper kinetochore corona may be required. We surmise that the achievement of “natural” recruitment conditions in our reconstitutions may promote an even more favorable reciprocal positioning of MAD1:MAD2 and BUB1:BUB3. Collectively, our observations are consistent with the idea that efficient SAC signaling requires the proximity and direct interaction of BUB1:BUB3 and MAD1:MAD2 in a single catalytic complex (Ballister et al., 2014; Heinrich et al., 2014; London and Biggins, 2014; Silio et al., 2015).

Crucially, BUB1:BUB3 becomes recruited to the kinetochore directly through an interaction with the MELT repeats of KNL1, triggered by their MPS1-dependent phosphorylation. Only few of the 19 MELT motifs of human KNL1 are occupied by BUB1:BUB3 during SAC activation (Jema et al., 2023; Krenn et al., 2014; Roy et al., 2020; Vleugel et al., 2015; Vleugel et al., 2013; Zhang et al., 2014). Engineered KNL1 constructs containing subsets of MELT repeats, including the single MELT1 with its neighboring KI and K2 motifs, or an array of six MELTs, were shown to be sufficient for a very robust SAC response (Jema et al., 2023; Krenn et al., 2014; Vleugel et al., 2015; Vleugel et al., 2013; Zhang et al., 2014). Thus, we believe that KNL1^Bonsai^, with its five MELT motifs including MELT1 (and associated KI1 and KI2 motifs) provides a sufficient number of MELT repeats to provide an “*in vivo*-like” response. Generating a full-length version of the 2316 (isoform 2, used here) or 2342-residue (isoform 2) KLN1 protein, which is mostly intrinsically disordered, turned out to be impractical.

Specific phosphorylation of MAD1 and BUB1 by MPS1 is essential to promote efficient MCC formation in our MCC assembly assay. Yet, we have no evidence of robust MPS1 recruitment to our recombinant kinetochores. We are currently unable to say whether this is due to the very rapid kinetics of MPS1 recruitment to kinetochores, which may prevent formation of a stable complex *in vitro*, or to the absence of the appropriate binding sites. The characterization of the binding sites for MPS1 may be incomplete. In unpublished results, we have observed normal MPS1 recruitment in the presence of several Ndc80C mutants originally proposed to impair kinetochore recruitment of MPS1 (Hiruma et al., 2015; Ji et al., 2015), possibly indicating that the determinants of this interaction remain to be identified.

Thus, an essential goal of our future studies will be to reproduce conditions for robust recruitment of all critical determinants of MCC assembly. Our *in vitro* reconstitution of the SAC monitors the time-dependence of MCC assembly in the absence of activities that disassemble MCC or that dephosphorylate the catalysts implicated in MCC assembly. Clearly, a fundamental function of kinetochores is to coordinate SAC signaling and microtubule binding, turning off MCC production upon biorientation. Such negative regulators include ATP-dependent, MCC-disassembly factors that counter the spontaneous assembly of MCC, phosphatases that remove crucial phosphorylation sites required for catalysis, and, in metazoans, molecular motors that disassemble the kinetochore corona upon microtubule binding (Fischer, 2023; Lara-Gonzalez et al., 2021b). The responsiveness of the SAC on kinetochore status depends on the function of these additional factors. Thus, another essential goal for future biochemical reconstitutions will be to include the negative regulators of MCC production.

## Materials and methods

### Production of recombinant proteins

Escherichia coli BL21 (DE3) cells containing vectors expressing KNL1M5 and its mutants were cultured in Terrific Broth at 37°C until reaching an OD600 of 0.8 - 1. At this point, 0.3 mM IPTG was added, and the culture was further incubated at 18°C for approximately 15 hours. Cell pellets were then suspended in lysis buffer (50 mM Hepes pH 8, 300 mM NaCl, 5% glycerol, and 2 mM TCEP) supplemented with a protease inhibitor cocktail (Serva). After sonication-induced lysis, the lysate was clarified by centrifugation at 90,000 g at 4°C for 1 hour. The resulting lysate was filtered (0.8 μm) and applied onto Ni Sepharose® High Performance beads that had been pre-equilibrated in lysis buffer. Following a 2-hour incubation at 4°C, the beads were washed with 50 volumes of lysis buffer and elution was performed with a buffer containing 300 mM imidazole. The eluted fractions containing the protein of interest were pooled, concentrated, and subjected to size exclusion chromatography (SEC). The SEC-purified fractions were concentrated, rapidly frozen in liquid nitrogen, and stored at -80°C. Escherichia coli BL21 (DE3) cells containing vectors expressing KNL1C underwent a similar cultivation process in Terrific Broth. After induction with 0.1 mM IPTG and subsequent incubation at 18°C for approximately 15 hours, cell pellets were lysed, clarified, and processed as described above. Cell pellets were suspended in lysis buffer (50 mM Tris pH 8, 50 mM NaCl, 5% glycerol, 2 mM TCEP). The cleared lysate was applied to Glutathione (GSH) beads, pre-equilibrated in lysis buffer and incubated at 4°C for 4 hours, allowing the GSH beads to selectively bind to the protein. The GSH beads were washed with lysis buffer and after extensive washing of these beads, the protein was cleaved overnight with a GST-Prescission protease to remove the GST tag and gain an untagged protein. The cleaved protein was then collected from the GSH beads and was concentrated using centrifugal filters with a 10 kDa mass cut-off (Merck). The concentrated protein sample was further purified using a Superdex 200 16/600 size-exclusion column, pre-equilibrated in SEC buffer (50 mM Hepes pH 8.0, 50 mM NaCl, 5 % glycerol, 2 mM TCEP). Other proteins were purified following detailed protocols (Faesen et al., 2017; Krenn et al., 2014; Pesenti et al., 2018; Petrovic et al., 2014; Piano et al., 2021).

### FRET measurements

The FRET sensors employed in this study were previously characterized in Faesen et al., 2017 and Piano et al., 2021. These sensors were utilized to investigate MCC assembly under various conditions. All measurements were conducted on a Clariostar plate-reader (BMG Labtech) using UV clear 96-well-plate flat-bottomed (Greiner), and data analysis and visualization were performed using Prism 9 software (Graphpad Software, Inc.). To monitor MCC assembly, the sensor components, along with MCC assembly catalysts, KMN, Rapamycin were combined in a final reaction volume of 100 μl, and the reading was initiated in buffer containing fresh 50 mM Hepes (pH 7.5), 150 mM NaCl, 5% glycerol, and 10 mM ß-Mercaptoethanol. Unless otherwise stated, assays utilized a final concentration of 100 nM for the FRET pair components (CFPBUBR1 & TAMRAMAD2) and were added the last before starting the measurements. The final concentration of catalysts varied depending on the specific experiment, which will be specified in the corresponding sections. Mixtures were excited using the filter 430-10 nm and the dichroic filter LP 504 nm, and the emissions were scanned from 450 to 650 nm. Single wavelength acceptor fluorescence measurements were performed using the emission filter 590-20 nm, reading every 90 seconds and mixing 60 seconds at 500 rpm just before starting the measurements; focal height 6 mm, 200 flashes, gain 1200.

### Analytical size-exclusion chromatography

All proteins were diluted to a concentration of 3 μM in 50 μl buffer (50 mM Hepes pH 8.0, 5% glycerol, 150 mM NaCl, 2 mM TCEP) and incubated for 1 hour at 4°C before loading on Superose 6 5/150 column (GE Healthcare) equilibrated with the same buffer on an AKTA-micro system (GE Healthcare). Elution of proteins was monitored at 280 nm. Fractions (100 μl) were collected and analyzed by SDS–PAGE and Coomassie blue staining.

### Pull-down experiments

For pulldown experiments, KNL1^C^ and KNL1^Bonsai^ is incubated together for ∼14 hours at 4°C with MPS1 kinase (1:10 kinase to substrate ratio) with 2 mM ATP, 10 nM MgCl_2_ to get phosphorylated KNL1^Bonsai^. To remove possible aggregates after the 14 hours incubation, overnight samples were then spun at max speed for 30 minutes in a benchtop centrifuge (at 4°C). Pre-equilibrated 10 µl dried GST beads for 1 hour on ice with 2 µM in 40 µl of GST-CENP-C^1-71^/GST-CENPC^1-544^. The beads were then transferred to Pierce micro-spin columns (Thermo Fischer Scientific). The supernatant was then removed by centrifugation (5 min, 1000 *g*, 4°C, same for all the following steps) and the remaining bait was added (KMN/CENP-11/CENP-11-KMN). After 1 hour incubation on ice, the supernatant was again removed and washed once before adding the prey. Again, after 1 hour incubation on ice, supernatant was removed. To remove the unbound material from the beads, beads were washed twice with 200 μl buffer (50 mM Hepes pH 8, 5% glycerol, 150 mM NaCl, 2 mM TCEP). The proteins were eluted in 25 μl buffer + 20 mM GSH pH 8, to which 5 μl 5x SDS-sample buffer were added.

### Cell culture

HeLa cells were grown in Dulbecco’s Modified Eagle’s Medium (DMEM; PAN Biotech) supplemented with 10 % tetracycline-free FBS (PAN Biotech) and 2 mM L-Glutamine (PAN Biotech). Cells were grown at 37°C with 5 % CO_2_.

### RNAi transfection

The following siRNAs oligos were used in this study: 60 nM siKNL1 (Invitrogen, HSS183683 5’-CACCCAGUGUCAUACAGCCAAUAUU-3’; HSS125942 5’-UCUACUGUGGUGGAGUUCUUGAUAA-3’; CCCUCUGGAGGAAUGGUCUAAUAAU-3’) for 24 h, 60 nM siZWINT (Sigma-Aldrich, 5’-GCACGUAGAGGCCAUCAAA-3’) for 48 h, 60 nM siNdc80C (Sigma-Aldrich, siNDC80^HEC1^ 5’GAGUAGAACUAGAAUGUGA-3’; siSpc24 5’-GGACACGACAGUCACAAUC-3’; siSpc25 5’-CUACAAGGAUUCCAUCAAA-3’) for 48 h.

siRNAs were transfected using RNAiMAX (Invitrogen) according to manufacturer’s instructions.

### Analysis of mitotic checkpoint functionality

10 000 HeLa cells were seeded in a 24-well imaging plate (ibidi) in DMEM and transfected with the respective siRNAs. After depletion, cells were synchronized in G2 phase with RO3306 (9 µM, Calbiochem). After 16 hours the treatment was washed out four times with pre-warmed PBS and media was replaced with CO_2_-independent L-15 imaging media supplemented with 10% FBS, 500 nM siR-DNA (Spirochrome) and 33 nM Nocodazole (Sigma-Aldrich). Cells were incubated for 6 hours before imaging. Cells were imaged at 37°C using a customized 3i Marianas system (spinning disk confocal) equipped with an Axio Observer Z1 microscope (Zeiss), a CSU-X1 confocal scanner unit (Yokogawa Electric Corporation, Tokyo, Japan), 40×/1.4 NA Oil Objective (Zeiss), and Orca Flash 4.0 sCMOS Camera (Hamamatsu). Images were acquired as z sections of 2 μm using Slidebook Software 6 (Intelligent Imaging Innovations). Manual count of mitotic cells was performed on maximal intensity projections using the software Fiji. Measurements were graphed with GraphPad Prism 10.2.

### Cell transfection and electroporation

For electroporation experiments, recombinant GFP-labelled KNL1^Bonsai^ was added to a final concentration 6 μM in the electroporation slurry, as previously described (Alex et al., 2019). A Neon Transfection System (Thermo Fisher Scientific) was used. Following an 8-hour recovery lag, cells were treated with 9 μM RO3306 (Calbiochem) for 15 hours. Subsequently, cells were released into mitosis in the presence of 3.3 μM Nocodazole (Sigma-Aldrich) for 1 hour before fixation for immunofluorescence or harvesting for immunoblotting.

### Immunofluorescence

Cells were grown on coverslips pre-coated with Poly-L-lysine (Sigma-Aldrich). For IF experiments cells were pre-permeabilized with 0.5% Triton X-100 solution in PHEM (Pipes, HEPES, EGTA, MgSO4) buffer supplemented with 100 nM microcystin for 5 mins and subsequently fixated with 4% PFA in PHEM for 20 mins.

Afterwards cells were blocked for 1 hour with 5% boiled goat serum (BGS) in PHEM buffer, and incubated for 2 hours at room temperature with the following primary antibodies: CREST/anti-centromere antibodies (Antibodies, Inc., 1:200) diluted in 2.5% BGS-PHEM supplemented with 0.1% Triton X100. Subsequently, cells were incubated for 1 hour sat room temperature with the following secondary antibodies: anti-goat anti-human Alexa Fluor 647 (Invitrogen, Carlsbad, California, USA). All washing steps were performed with PHEM-T buffer. DNA was stained with 0.5 lg/ml DAPI (Serva), and Mowiol (Calbiochem) was used as mounting media.

### Cell imaging

Cells were imaged at room temperature using a spinning-disk confocal device on the 3i Marianas system equipped with an Axio Observer Z1 microscope (Zeiss), a CSU-X1 confocal scanner unit (Yokogawa Electric Corporation), 100 × /1.4NA Oil Objectives (Zeiss), and Orca Flash 4.0 sCMOS Camera (Hamamatsu). Images were acquired as z sections at 0.27 μm (using Slidebook Software 6 from Intelligent Imaging Innovations or using LCS 3D software from Leica). Images were converted into maximal intensity projections, exported, and converted into 16-bit TIFF files. Figures were arranged using Adobe Illustrator 2022.

### Immunoblotting

Mitotic cells were collected through shake-off and resuspended in lysis buffer (150 mM KCl, 75 mM Hepes [pH 7.5], 1.5 mM EGTA, 1.5 mM MgCl_2_, 10% glycerol, and 0.075% NP-40 supplemented with protease inhibitor cocktail [Serva] and PhosSTOP phosphatase inhibitors [Roche]). After lysis the whole-cell lysates were centrifuged at 15,000 rpm for 30 mins at 4°C.

Afterwards, the supernatant was collected and resuspended in sample buffer for analysis by SDS-PAGE and Western blotting. The following primary antibodies were used: GFP (rabbit, in-house, 1:1,000) and Tubulin (mouse monoclonal, 1:8,000; Sigma-Aldrich). As secondary antibodies, anti-mouse or anti-rabbit (1:10,000; NXA931 and NA934; Amersham) conjugated to horseradish peroxidase were used. After incubation with ECL Western blotting reagent (GE Healthcare), images were acquired with the ChemiDoc MP System (Bio-Rad) using Image Lab 6.0.1 software.

## Acknowledgements

We thank Andrea Ciliberto for critical reading of the manuscript, all members of the Musacchio laboratory for helpful discussions, Amal Alex, Sara Carmignani, Carolin Körner, Sabine Wohlgemuth, Ingrid Hoffmann, Isidora Arias, and Javier Aviles for help with reagents preparation. A.M. acknowledges funding from the Max Planck Society, the European Research Council (ERC) Synergy Grant 951430 (BIOMECANET), the DGF’s Collaborative Research Centre 1430 “Molecular Mechanisms of Cell State Transitions”, and the CANTAR network under the Netzwerke-NRW program.

## Author contributions

**Conceptualization:** S.S., V.P, A.M.

**Investigation:** S.S., V.P., V.C., S.G.

**Funding acquisition:** A.M.

**Project Administration:** A.M.

**Resources:** P.S.

**Supervision:** A.M.

**Validation:** S.S., V.P, A.M.

**Visualization:** S.S., A.M.

**Writing – original draft:** A.M.

**Writing – review & editing:** All authors

## Supplementary figures

**Figure S1.**
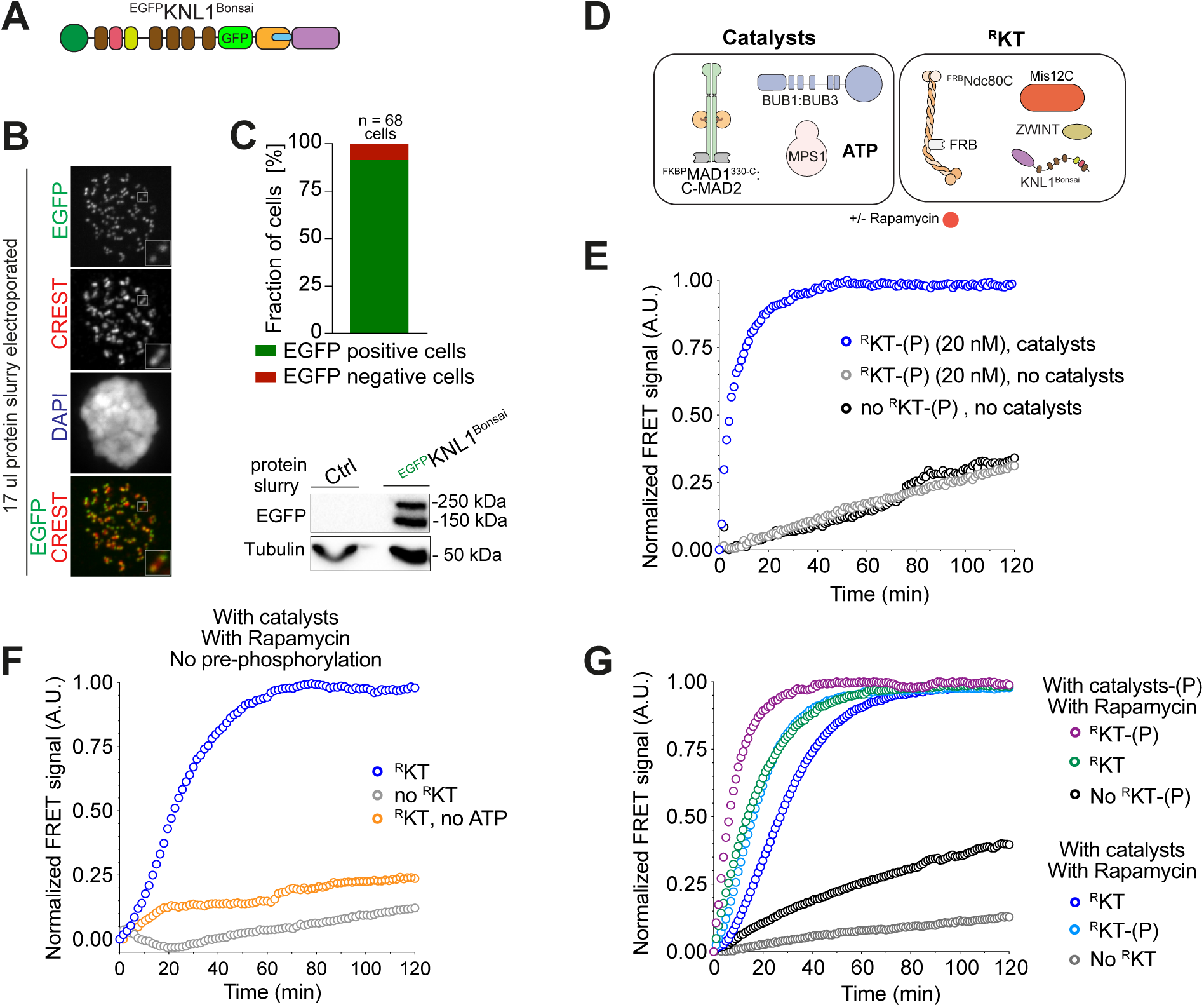
Further characterization of ^R^KTs. **A)** Scheme of EGFP-labelled KNL1^Bonsai^ used in panel B. **B)** Representative image of HeLa cells electroporated with ^GFP^KNL1^Bonsai^ (at a slurry concentration of 6 µM). DAPI and CREST were used to stain chromosome DNA and inner kinetochores, respectively. Scale bar: 5 µM. **C)** Quantification of the electroporation efficiency with the immunoblot from mitotic lysates obtained from HeLa cells electroporated with ^GFP^KNL1^Bonsai^. 50 μg cleared lysate was used, and tubulin was used as a loading control. The two bands in the anti-GFP blot represent ^EGFP^KNL1^Bonsai^ and unfused ^EGFP^KNL1^M5^. **D)** Scheme of proteins used in assays in panels E-F. **E)** Sensitized fluorescence emission (FRET) curves for MCC sensor in presence of catalysts and kinetochore (blue, same as shown in Figure 3B) no kinetochores (red, same as shown in Figure 3B), or in two conditions where catalysts were omitted (black and grey). Catalytic components or kinetochores were phosphorylated with MPS1 prior the assay as indicated. The blue and red curve were normalized to their own maximum value (already shown in Figure 3B). The grey and black curve were normalized to the maximum value of the blue curve. **F)** FRET assay monitoring the rate of MCC formation in the presence (blue curve, already shown in Figure 3D) or the absence (grey curve; already shown in Figure 3D) of reconstituted ^R^KTs, and in the absence of ATP (orange). MAD1:MAD2, BUB1:BUB3 and ^R^KT were not pre-phosphorylated with MPS1. **G)** FRET assay monitoring the rate of MCC formation with MPS1-pre-phosphorylated catalysts (20 nM MAD1:MAD2 and 40 nM BUB1:BUB3; purple, green black curves) or not pre-phosphorylated catalysts (blue, light blue, grey curves). ^R^KT in pre-phosphorylated or not pre-phosphorylated forms were added or omitted as indicated. Purple, green, light blue and blue curves were normalized to their own maximum value. The black and grey curve were normalised to the maximum value of green curve.

**Figure S2.**
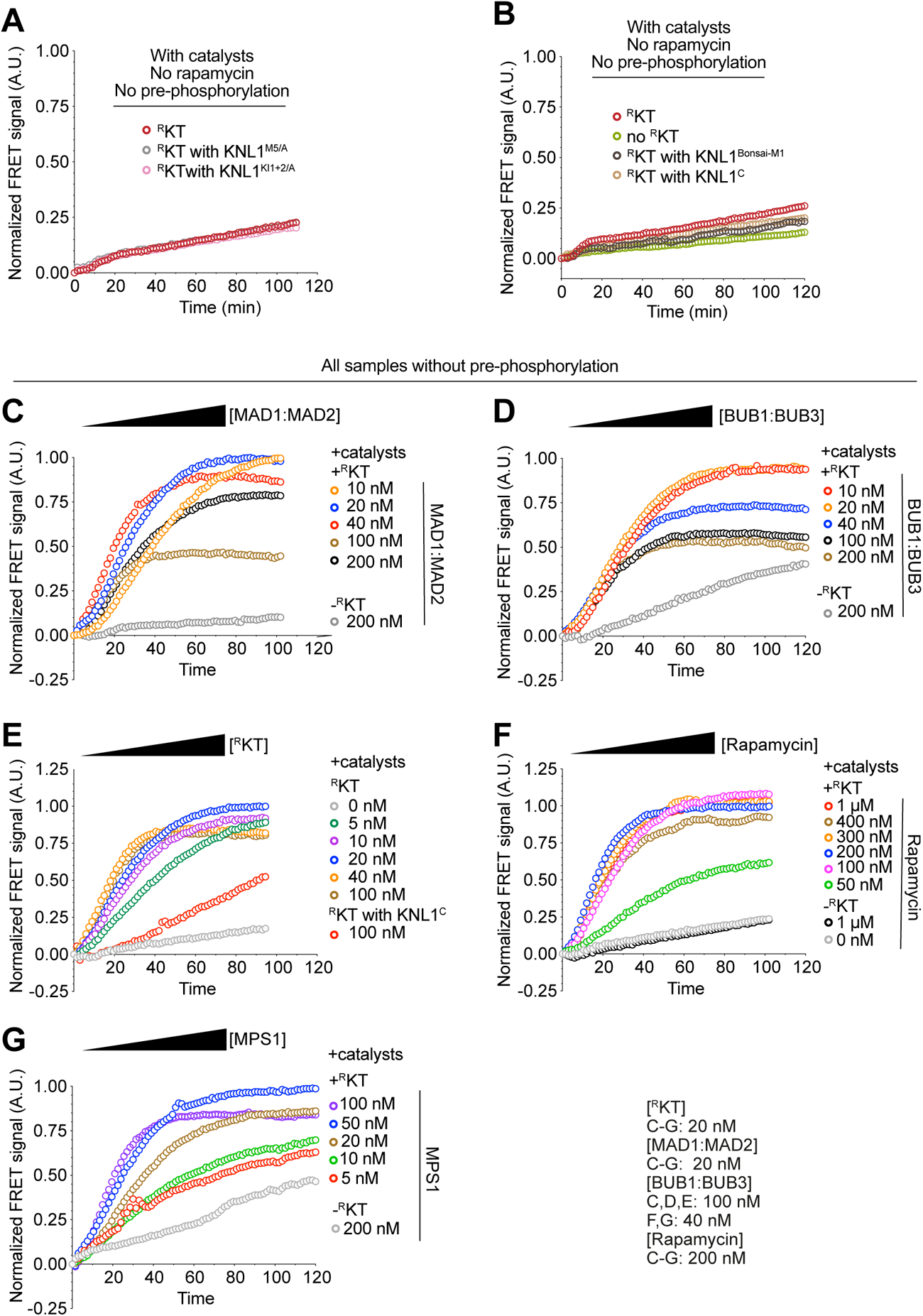
Rate of MCC assembly under various conditions. **A)** Additional controls for experiments with ^R^KT performed in Figure 4C. FRET assay comparing the MCC catalytic assembly with wild-type ^R^KT (maroon), ^R^KT with Knl1^KI1+2/A^ (where KI1 and KI2 motifs of KNL1 are mutated, pink) and ^R^KT with Knl1^M5-A^ (where five MELT motifs of KNL1 are mutated, grey) without Rapamycin. **B)** Additional controls for experiments with ^R^KT performed in Figure 4D. FRET assay comparing the MCC catalytic assembly with wild-type ^R^KT (maroon), ^R^KT with Knl1^M1/A^ (where only first MELT motif is present, dark grey), ^R^KT with Knl1^C^ (where only C terminal of KNL1 is present light brown), and no ^R^KT (light green) in the absence of Rapamycin. **C**-**G**) Effect of titration of the indicated catalyst concentration, Rapamycin, and ^R^KT on MCC assembly rate.

**Figure S3.**
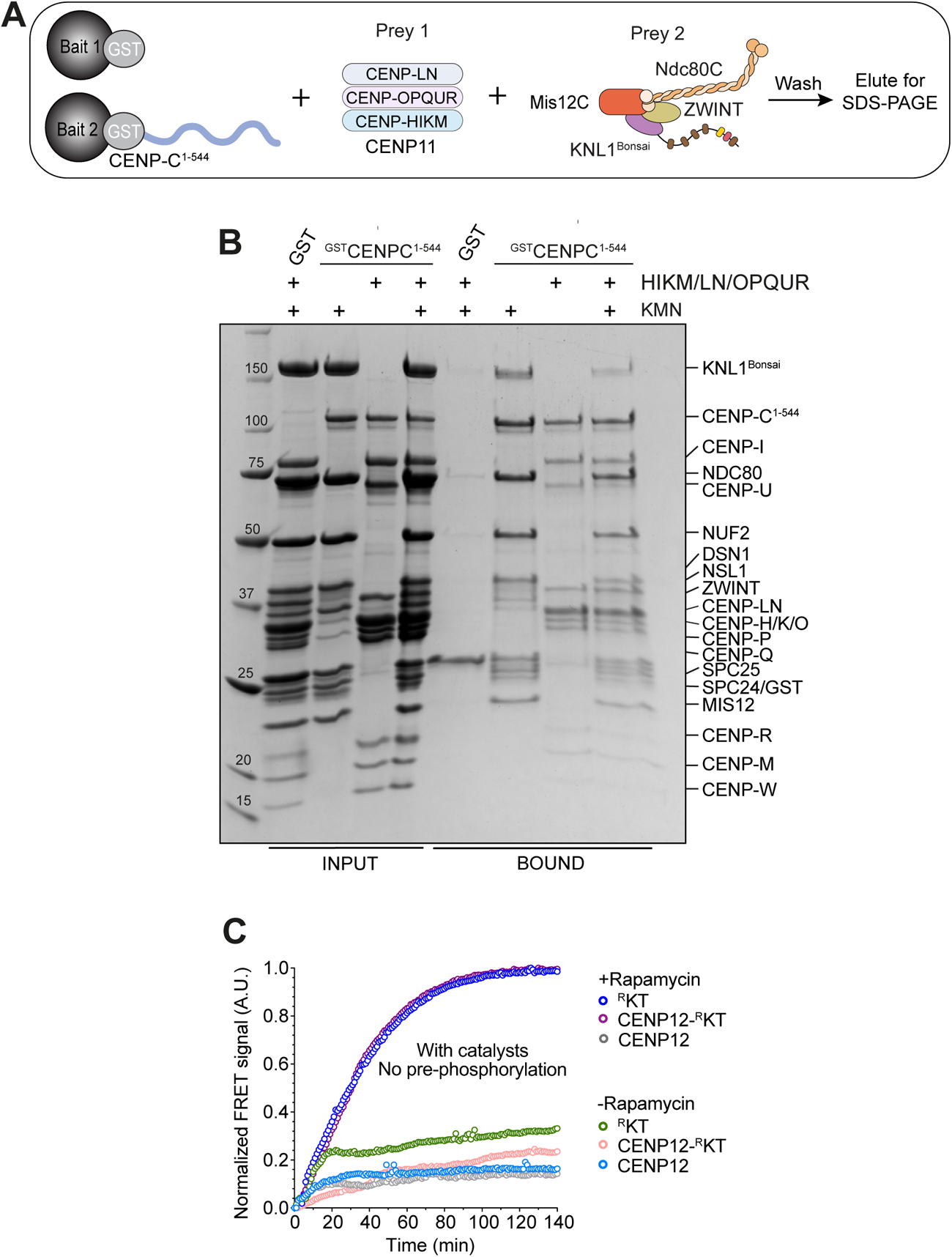
CCAN does not contribute to catalytic assembly of MCC. **A)** Scheme of GST pulldown assays. GST or GST-CENP-C were incubated with CENP12 on GSH beads as bait. ^R^KT was added as the prey **B)** SDS-PAGE of GSH based pulldown with the indicated baits and preys. GST-CENP-C^1-544^ (3 μM) as a bait with CENP-12(HIKM/LN/OPQUR) (4 μM) and ^R^KT (6 μM) as prey **C)** FRET assay monitoring the rate of MCC formation in the presence of the indicated CCAN species and in the presence of catalysts and ^R^KT. The blue and purple curves were normalized to their maximum. Fluorescence values of the other curves were normalized to the maximum value of the blue curve..

**Figure S4.**
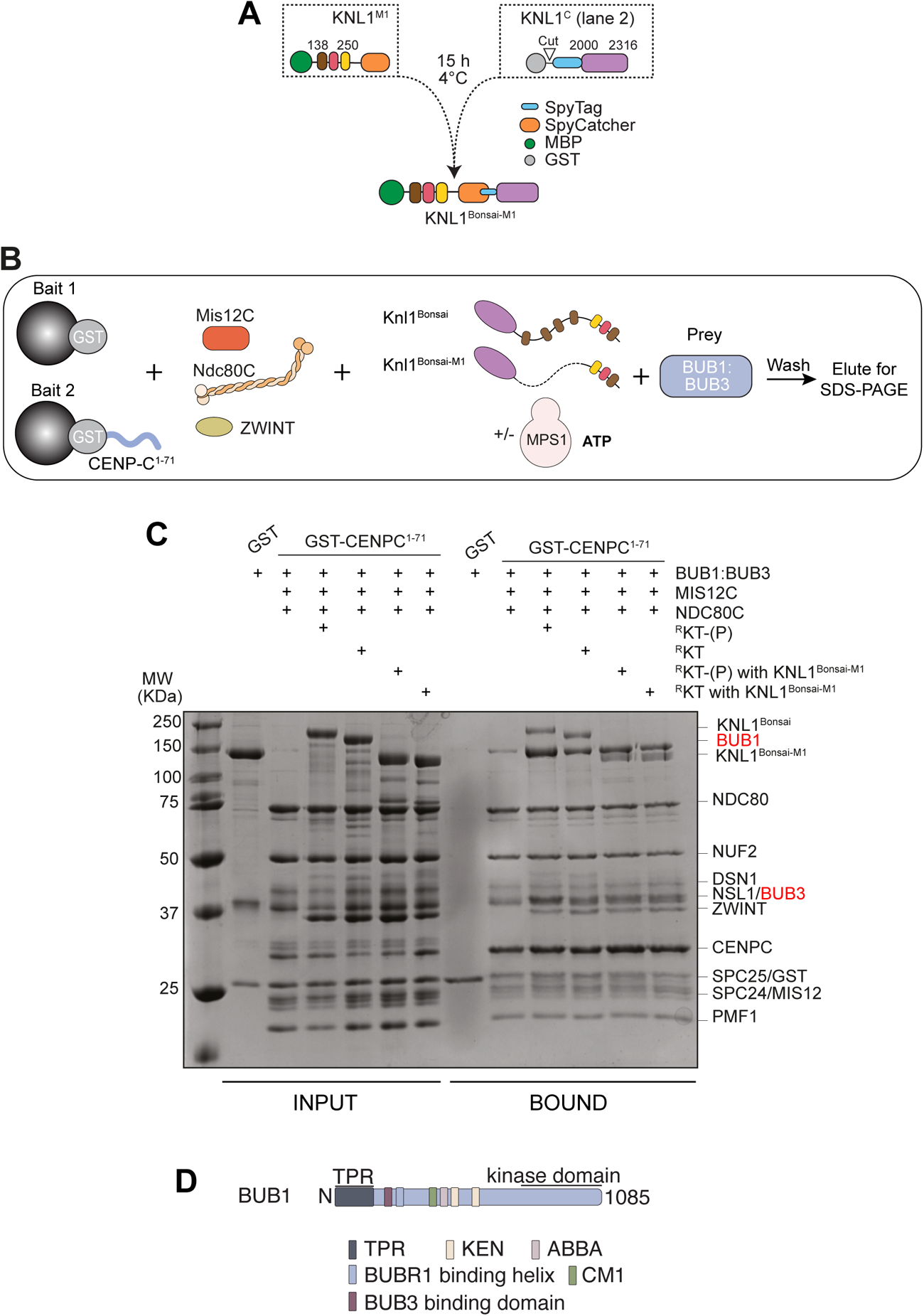
Additional experiments with KNL1 and BUB1 variants. **A)** Scheme for the assembly of KNL1^Bonsai-M1^. **B)** Scheme for binding assays with ^R^KT assembled with KNL1^Bonsai^ or KNL1^Bonsai-M1^. **C)** SDS-PAGE of BUB1 binding assays with ^R^KT assembled with KNL1^Bonsai^ or KNL1^Bonsai-M1^. **D)** Scheme of BUB1 motifs and domains

